# Fragile polyQ assemblies cause Golgipathy in Huntington’s disease

**DOI:** 10.1101/2024.10.06.616845

**Authors:** Lixiang Ma, Xinyu Chen, Yang Liu, Lijun Dai, Fangzhou Ye, Weiqi Yang, Hada Buhe, Jixin Ma, Chenyun Song, Li Li, Dandan Fan, Fanxun Chen, Haoman Chen, Jianwei Shuai, Jianzhong Su, Hexige Saiyin

## Abstract

Huntingtin (HTT) is a naturally aggregating protein that causes Huntington’s disease (HD) when its polyglutamine (polyQ) tract exceeds 38 repeats. Despite its importance, the biology of HTT aggregates remains poorly defined. Here, using high-resolution imaging of cells derived from HD families, we redefine polyQ assemblies, traditionally regarded as pathogenic aggregates, as dynamic and structurally organized intracellular compartments resembling knitted-fabric patches. These assemblies closely associate with and encircle the Golgi apparatus, integrating Golgi ribbons and stacks to form a previously unrecognized polyQ assembly–Golgi complex. Mechanistically, fragmentation of polyQ assemblies is dynamically coupled with mitotic Golgi fragmentation, whereas inhibition of ADP-ribosylation factor (ARF) function by Brefeldin A disrupts and fragments the complex, revealing an intimate structural and functional coupling between polyQ assemblies and Golgi architecture. The presence of mutant HTT (mHTT) destabilizes these assemblies and complexes, altering their response to nutrient deprivation and autophagy enhancers but not to antisense oligonucleotide (ASO) therapy. Functionally, polyQ assemblies in HD cells compromise the scaffolding capacity of the Golgi apparatus, clathrin-coated vesicles, and ARF1, impair Golgi glycosylation, and promote aberrant neuronal firing activity. Collectively, these findings establish polyQ assemblies as dynamic structural regulators of Golgi organization and function and demonstrate that mHTT disrupts the homeostatic dynamics of the polyQ assembly–Golgi complex, leading to Golgipathy.

## Introduction

Huntington’s disease (HD) is a devastating, progressive neurodegenerative disorder histopathologically characterized by irreversible neuronal loss within the striatum, which manifests clinically as profound motor dysfunction and cognitive decline (Kernich, 2002; Saudou and Humbert, 2016; Walker, 2007a). HD is driven by an abnormal expansion of a polyglutamine (polyQ) tract within the huntingtin (HTT) protein, a mutation that renders the protein highly prone to misfolding and macromolecular aggregation (Gutekunst et al., 1999). Notably, HTT aggregates exhibit significant instability following chemical fixation (Ferrante et al., 1997). This persistent histological challenge can inadvertently introduce structural artifacts, potentially leading to the misinterpretation of polyQ biology and pathophysiology if standardized sample preparation protocols are not strictly enforced.

Although the formation of polyQ aggregates and the deposition of macromolecular intranuclear inclusion bodies are universally recognized as pathological hallmarks of HD (DiFiglia et al., 1997; Gutekunst et al., 1999), their precise role remains controversial. Emerging evidence suggests a paradox: some studies indicate that aggregate formation may actually increase single-neuron survival (Arrasate et al., 2004), exerting a protective rather than toxic effect (Kim et al., 1999; Saudou et al., 1998). Conversely, soluble monomeric HTT and its transition into oligomeric species, which precede fibril and inclusion body formation, are the primary drivers of cellular toxicity (Hatters et al., 2013; Jimenez-Sanchez et al., 2017; Lajoie and Snapp, 2010; Legleiter et al., 2010; Mukai et al., 2005; Poirier et al., 2002; Takahashi et al., 2008). These observations suggest that mature HTT aggregates may possess critical biological functions rather than being purely pathogenic.

HTT is a stiff protein composed of multiple ɑ-helical and polyQ regions (Dougan et al., 2009; Guo et al., 2018; Saudou and Humbert, 2016; Sivaramakrishnan et al., 2008; Tabrizi et al., 2020). Expanded polyQ (>35) and short polyQ form a rigid fibril (Bauerlein et al., 2017; Dougan et al., 2009; Isas et al., 2017; Scherzinger et al., 1999); however, expanded polyQ fibrils have been reported to physically disrupt organelle membranes, such as the endoplasmic reticulum (ER), and alter membrane dynamics (Bauerlein et al., 2017). Functionally, HTT acts as a versatile scaffold involved in a wide array of cellular processes, including vesicular transport, autophagy, gene transcription, and embryonic development (Saudou and Humbert, 2016). HTTs are essential proteins in embryonic development (Dragatsis et al., 1998; Duyao et al., 1995; Zeitlin et al., 1995). The presence of mutant HTT (mHTT) precipitates widespread cellular dysfunction, ranging from early transcriptional dysregulation and synaptic impairment to mitochondrial distress and disruption of the nuclear pore complex (Costa et al., 2010; del Toro et al., 2009; DiFiglia et al., 1997; Grima et al., 2017; Kim et al., 2010; Mehta et al., 2018; Milnerwood et al., 2006; Saudou and Humbert, 2016; Shirasaki et al., 2012). mHTT induces extrasynaptic excitotoxicity (Milnerwood et al., 2010), higher levels of brain inflammation (Lee et al., 2020; Silvestroni et al., 2009), abnormal fetal brain development (Barnat et al., 2020), and aging (Machiela and Southwell, 2020). HAP40 stabilizes HTT(Guo et al., 2018; Harding et al., 2021). HTT interacts with and activates dynamin, a protein that catalyzes membrane fission (Caviston et al., 2007; Caviston et al., 2011; Morlot and Roux, 2013), microtubule motors, and actin-associated adaptors to support cellular vesicle transport, including autophagy, endosomes, and lysosomes (Caviston et al., 2011; Olenick and Holzbaur, 2019).

Using HD family-derived cells and super-resolution scanning, we revealed a knitted-fabric-like network formed by HTT polyQ that encircles flat Golgi stacks/cisternae and dynamically couples with both mitotic and stress-related Golgi fragmentation. We demonstrate that while normal polyQ assemblies support the structural integrity of the Golgi, pathogenic mHTT crisps these assemblies, destabilizing the complex and impairing Golgi activities. Our findings reveal that the polyQ assembly-Golgi complex is a highly organized functional unit, the disruption of which represents a central mechanism of mHTT-induced toxicity.

## Results

### Characterization of endogenous polyQ aggregate structure

We identified an HD family in the Mongolian population of West China. 24 persons have identifiable syndromes, including an adult-onset, abnormal gait, uncontrolled arm, leg, and head movements, and psychiatric abnormalities; some patients with severe syndromes showed mania and pica **(Table S1).** The self-reported symptoms appeared at ages 24 to 50; patients died between 35 and 56 **(Table S1)**. Genetic testing revealed that the patients in this family harbor 44 to 59 CAG repeats in *HTT* **(Fig. 1A, B; Table S1)**, compared with 18-19 CAG repeats in healthy individuals. Consistent with HD, the lengths of CAG repeats define the disease onset age, symptom severity, and death age **(Fig. 1C; Table S1).**

**Figure 1.**
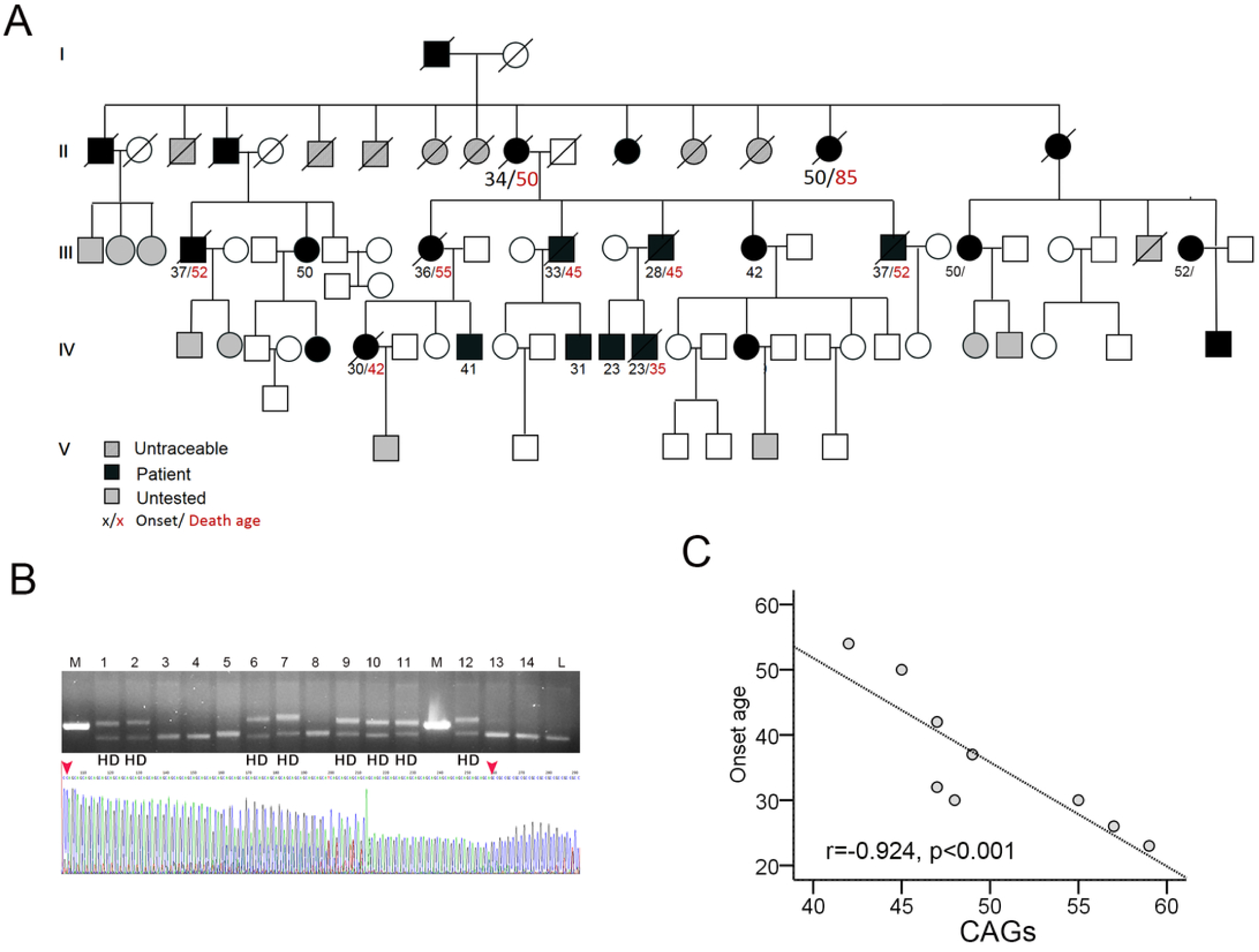
Identification of a family with HD. **A.** The pedigree of the family with HD. **B.** Representative PCR and Sanger sequencing analysis of HTT CAG repeat lengths. Upper panel: PCR amplification shows that 8 individuals (8/14) contain the expanded allele; the upper band represents the expanded CAG repeat tract. Lower panel: Sanger sequencing chromatogram confirms the presence of expanded CAG repeats in an individual with the expanded allele (49 repeats; red arrows denote the boundaries of the CAG repeat region). M, positive control allele (35 CAG repeats); L, a healthy local man. See also Table S1. **C.** Spearman’s rank correlation analysis between CAG repeat length and the age of disease onset.

To assess polyQ status in the fibroblasts of this HD family, cells were stained with the 3B5H10 antibody, which recognizes both HTT and mHTT polyQ stretches with equal affinity (Owens et al., 2015). We observed that the polyQ structures in both patient and healthy sibling fibroblasts organized into distinctive, fabric-like assemblies. These structures resembled knitted-fabric patches formed by parallel spindles that merged at regular intervals, mostly taking on a chain-of-rings appearance (**Fig. 2A)**. Contrary to other observations (Tousley et al., 2019), phalloidin counterstaining showed that the polyQ assembly did not regularly parallel actin filaments and randomly interacted with actin filaments **(Fig. 2A)**. 3D rendered tomography showed that the polyQ assembly in fibroblasts interacts with the nuclear membrane **(Fig. 2A)**. Notably, the size of polyQ assemblies in HD is similar to that in healthy fibroblasts (HD: n=4; healthy: n=4) **(Fig. 2B, C)**. Mitochondria did not preferentially enrich at the polyQ assembly site: some of them were present in the polyQ assembly network’s space, while others were not **(Fig. 2D).** Over 300 proteins in humans have polyQ repeats that exceed 7(Mier and Andrade-Navarro, 2021). 3B5H10 antibody was reported to bind other polyQ-containing proteins, including androgen receptor, atrophin, and ataxin-3 (Miller et al., 2011). To see whether the polyQ assemblies detected by the 3B5H10 antibody contain full-length or large fragments of huntingtin (HTT), we selected two distinct HTT antibodies: EM48, which binds the first 256 amino acids (excluding the polyQ stretch), and 3E10, which targets the HDA region (amino acids 1171–1177). In human fibroblasts, the immunostaining patterns for both EM48 and 3E10 were nearly identical to those observed with 3B5H10, and closely circled the GM130-stained Golgi apparatus **(Fig.S1A, B).** To see the C-terminal position of HTTs in polyQ assembly, we have stained the polyQ with a 3B5H10 antibody and the C-terminal with a C-terminal-specific antibody (LSB11011) and found that discontinuous C-terminal antibody signals colocalize with the polyQ assemblies **(Fig. 2E)**. Notably, we observed that a significant fraction of full-length HTT, marked by both C- and N-terminal antibodies, docked onto actin filaments through the N-terminus **(Fig.S1-D).** This population is present on the ruffling membrane, membrane curvature, large vacuoles, and the edge of large polyQ assembly **(Fig.S1C-D)**, indicating that the spatial position of an HTT in large polyQ assembly is unique. The unique spatial arrangement of HTT in large assemblies might hide the C-terminus and restrict the binding of the C-terminal antibody to its epitope. The polyQ assemblies were fragmented in mitotic cells **(Fig. 2F).** The polyQ fragments in the mitotic cells were labeled by the C-terminal antibody **(Fig. 2F).**

**Figure 2.**
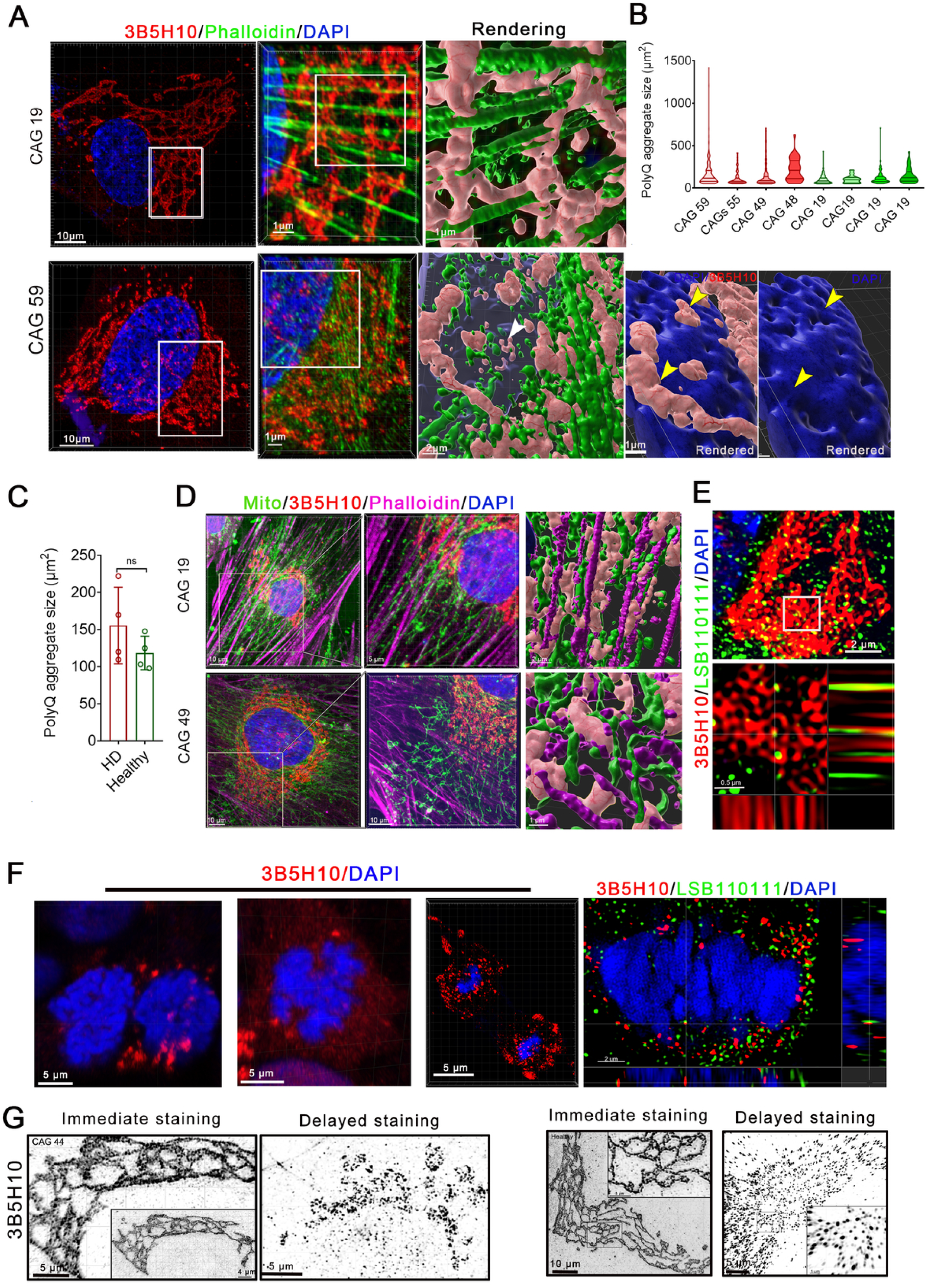
Morphological characterization of endogenous polyQ. **A.** Representative confocal immunofluorescence images of healthy control (19 CAG repeats) and HD patient (59 CAG repeats) fibroblasts stained with the 3B5H10 antibody and phalloidin. Right panels show 3D tomographic reconstructions generated via IMARIS software, illustrating the spatial distribution of polyQ assemblies relative to the nuclear surface and F-actin filaments (white arrows, intranuclear polyQ puncta; yellow arrows, polyQ assemblies interacting with the nuclear envelope/DAPI-rendered surface). **B.** The size of polyQ assemblies in the fibroblasts of healthy siblings (n=4; cells, n=306) and HD patients (n=4; CAGs: 59, 55, 49, and 47; cells, n=783). Fiji ImageJ was used to measure the assembly size. **C.** Measurements of the average size of polyQ assemblies in HD patient and healthy sibling fibroblasts. Data are presented as mean ± SEM. Statistical significance was evaluated using a two-tailed Student’s t-test; ns, p > 0.05. **D.** Mito-tracker, phalloidin, and 3B5H10 antibody immunostaining in the fibroblasts of HD patients (CAG, 49) and healthy siblings. 3D rendering tomography in the right two panels revealed the spatial relationships of polyQ assemblies with mitochondria and actin filaments. **E.** Representative immunofluorescence images of HD fibroblasts co-stained with an HTT C-terminal-specific antibody (LSB11011) and an anti-polyQ antibody (3B5H10), demonstrating the colocalization of C-terminal huntingtin fragments within cytoplasmic polyQ assemblies. The lower sectional view of the upper 3D image is from the white rectangle. **F.** 3B5H10 (three right) or 3B5H10 (N-terminal) and LSB11011 (C-terminal) (left) stained polyQ puncta/patches in mitotic cells. **G.** Representative images of 3B5H10 antibody immunostaining in HD patient (CAG 44) and healthy sibling fibroblasts by two different protocols (immediate staining is performed 15-30 minutes after 4% PFA fixation; delayed staining is performed 48 h after 4% PFA fixation). Left panels show freshly fixed and immediately stained groups; right panels show freshly fixed and later stained groups. Images were taken by SIM microscopy with Z-stacks.

The HTT fragment of the first exon is pathogenic and widely used for constructing HD models (Brooks and Dunnett, 2015; Yang et al., 2020). To see if the transfected exogenous HTT fragment of the first exon can be recruited into polyQ assemblies, we transfected the fibroblasts with three vectors of the *HTT* first exon containing 19, 23, and 74 CAGs, respectively. The transfected first exon fragment of *HTT* rarely localizes to and is recruited into the Golgi apparatus **(Fig. S1F, G).** Strikingly, the polyQ assemblies fragmented after 4% PFA fixation: the immediate staining after fixation is the only way to visualize an intact polyQ assembly **(Fig. 2G)**. These findings indicate that polyQ assemblies are unique, dynamic polyQ assemblies with a specific structure in cells.

### PolyQ assemblies of HD patients and healthy individuals differentially respond to stress and drugs

Antisense oligonucleotides (ASOs) reduce the transcription of *HTT (Kordasiewicz et al., 2012)*. To test how the polyQ assemblies in fibroblasts respond to the ASO, we treated the HD family fibroblasts with ASO (20 and 40 µM) and found that the ASO equally reduced the polyQ assemblies in healthy and HD fibroblasts without affecting the structure of polyQ assembly **(Fig. 3A-C).** Autophagy-enhancing agents were reported to reduce mHTT (Martin et al., 2015). We also treated HD fibroblasts with onjisaponin F (5, 10, 20, and 40 µM), an active component of Polygala tenuifolia (Nagai et al., 2001). Onjisaponin F reduced polyQ assemblies in HD fibroblasts, but not those in healthy fibroblasts, in a dose-dependent manner, without fragmenting polyQ assemblies **(Fig. 3D, E)**. To see how the polyQ assemblies respond to energy deprivation, we cultured the fibroblasts from 4 patients with HD (CAG: 48, 49, 55, and 59) and 2 healthy siblings in a low-glucose medium. We fixed cells at 24, 48, and 72 h and evaluated the polyQ assemblies. Under glucose deprivation, the polyQ assemblies in HD fibroblasts are diminished and fragmented over time. In contrast, polyQ assemblies in healthy fibroblasts are stable, and the smaller polyQ fragments in healthy cells are decreased **(Fig. 3F-H)**. After culturing in the low-glucose medium for 72 h, we switched to a high-glucose medium. After culturing for an additional 48 h, we observed that the polyQ assemblies in HD fibroblasts had been restored **(Fig. 3F-H)**. Together, the presence of pathogenic polyQ shifts the response of polyQ assemblies to energy deprivation and to an autophagy-enhancing treatment in fibroblasts.

**Figure 3.**
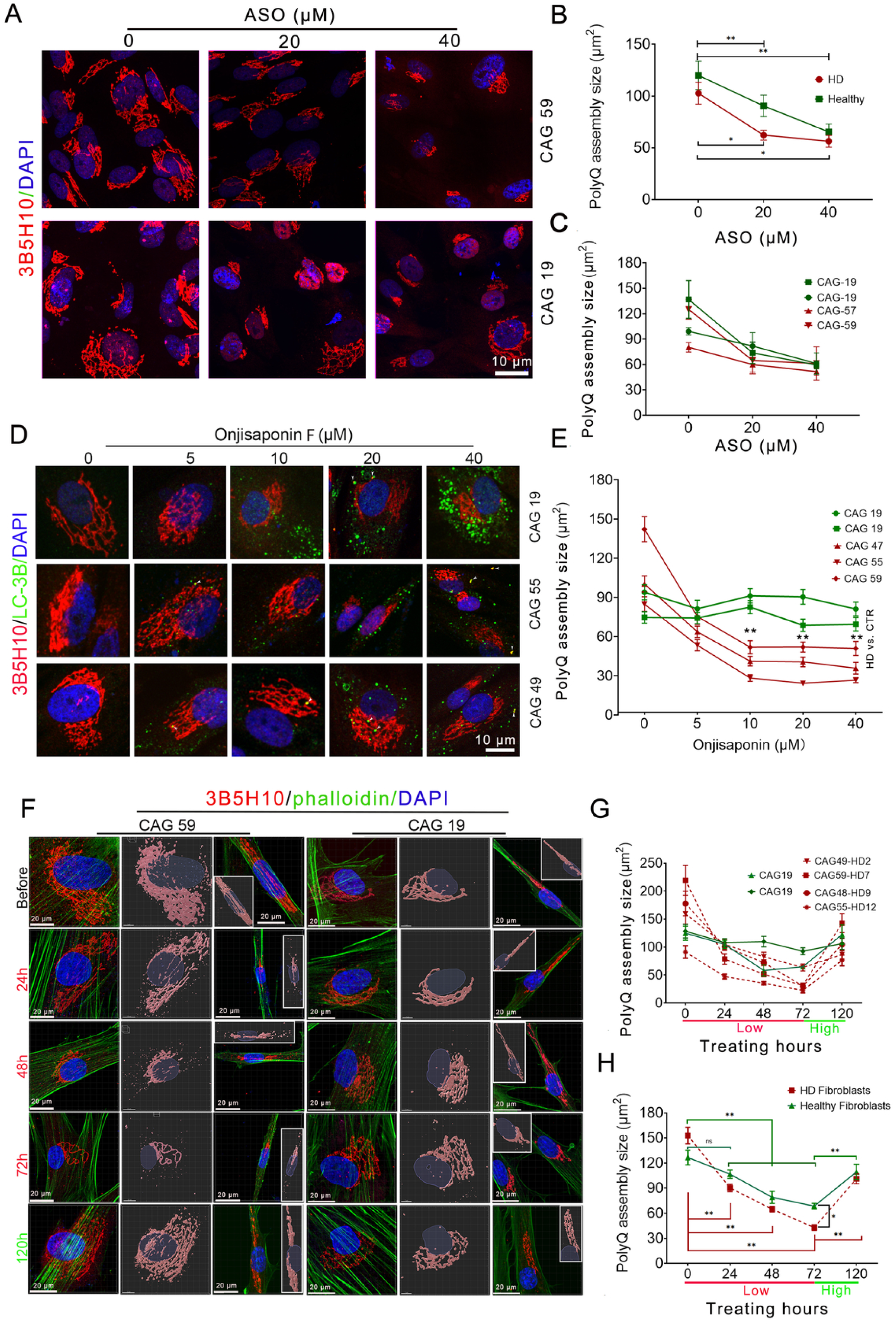
PolyQ assemblies of HD and healthy individuals differentially respond to stress and pharmacological treatments. **A.** Representative immunofluorescence images of polyQ assemblies in antisense oligonucleotide (ASO)-treated HD and healthy fibroblasts, stained with the 3B5H10 antibody. **B, C.** Measurements of changes in polyQ assembly size following ASO treatment. Comparison of assembly size trends between HD and healthy individual cohorts (B). Assembly size changes tracked within individual cell lines (C) (HD, n = 2 [CAG 57 and 59]; Healthy, n = 2; Two-way ANOVA, \**p* < 0.05, \*\**p* < 0.01). **D.** Immunostaining of Onjisaponin F-treated HD and healthy fibroblasts co-labeled with LC-3B and 3B5H10 antibodies. **E.** Measurements of polyQ assembly size changes following Onjisaponin F treatment in HD and healthy individuals (HD, n = 3; Healthy, n = 2; Two-way ANOVA, \*\**p* < 0.01). **F.** 3B5H10-stained polyQ assemblies under glucose starvation in the HD and healthy fibroblasts at different time points. The middle panels and the inserted image (second panel) are 3D rendering tomography in IMARIS. **G, H.** Temporal changes in polyQ assembly size under glucose starvation in fibroblasts from HD patients and healthy siblings. Assembly size changes tracked within individual cell lines (red lines: HD patients; green lines: healthy individuals) (G). Cohort-level comparison of polyQ assembly size changes between HD and healthy groups(H) (Two-way ANOVA, \*\**p* < 0.01).

### PolyQ assemblies circle the Golgi stack and couple to Golgi dynamics

HTTs are present on the surface of the Golgi, and mHTT impairs Golgi trafficking (del Toro et al., 2009; del Toro et al., 2006; DiFiglia et al., 1995). To our knowledge, the appearance of polyQ assemblies resembles Golgi ribbons. GM130 antibody, a Golgi marker, and 3B5H10 antibody coimmunostaining revealed that polyQ assemblies circle flat Golgi ribbon/stacks; the flat Golgi stack stands between the internal space of two parallel spindles of polyQ assemblies, and an isthmus-like hollow in the Golgi appeared at where polyQ circled the Golgi **(Fig. 4A)**, indicating the physical interactions of the Golgi apparatus with polyQ assemblies. The size of the Golgi apparatus in HD fibroblasts is smaller than that of healthy cells **(Fig. 4B).** Clathrin-coated vesicles sort at the Golgi (Klumperman, 2011). Clathrin+ is preferentially distributed along or docked with the side of polyQ assemblies opposite to the F-actin bundle **(Fig. 4C, D),** and polyQ assemblies in HD fibroblasts attached fewer clathrin^+^ vesicles than polyQ assemblies in healthy fibroblasts **(Fig. 4E).** Huntingtin-associated protein 40 (HAP40), a protein coevolved with HTT, binds to all three domains of HTT and stabilizes HTT (Guo et al., 2018; Harding et al., 2021). PolyQ assemblies in HD and healthy cells attached the same amount of HAP40, which resided on the corners of polyQ assemblies **(Fig. 4F, G).**

**Figure 4.**
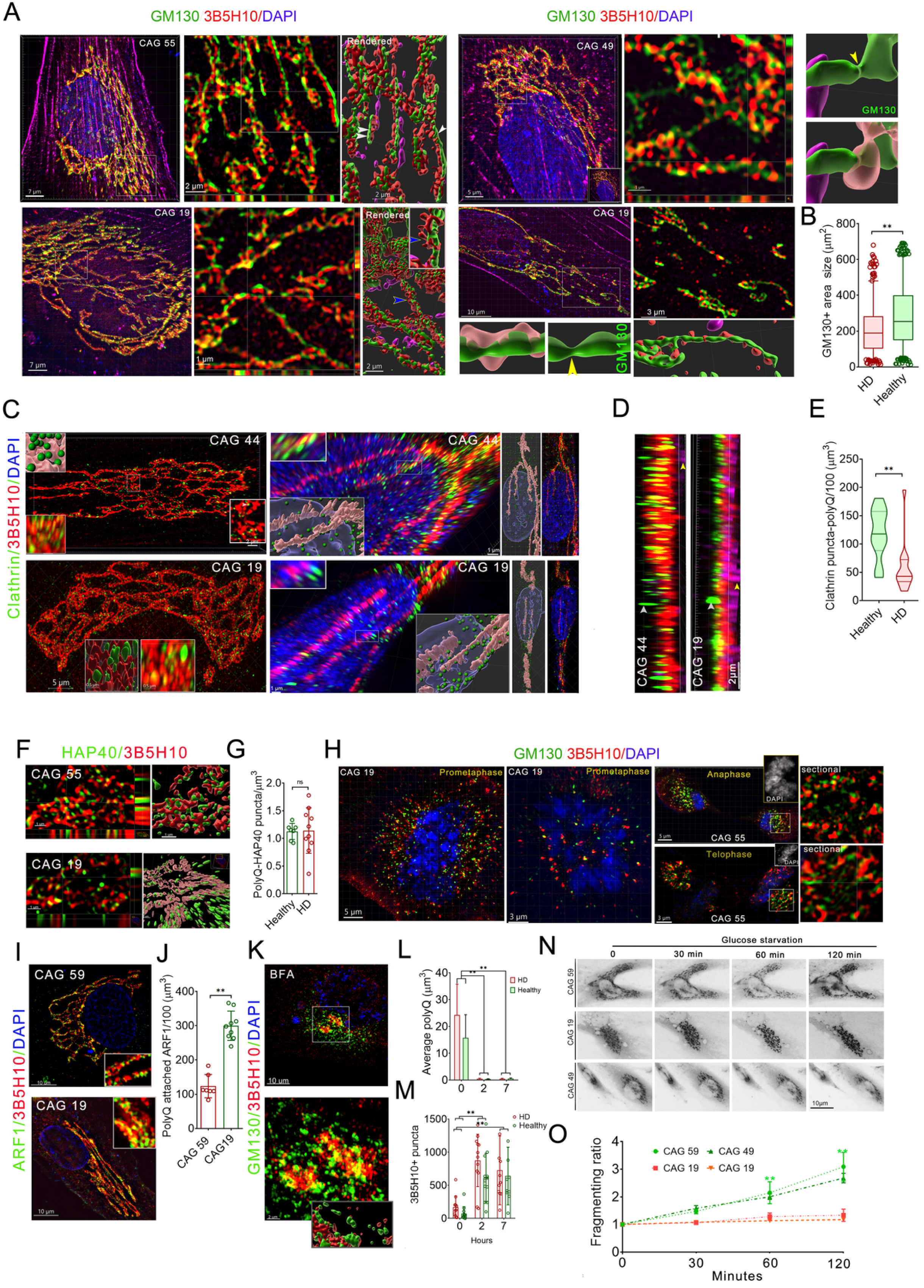
PolyQ assemblies circle the large Golgi apparatus. **A.** Immunofluorescence staining of GM130, 3B5H10, and phalloidin in healthy sibling and HD fibroblasts demonstrates that polyQ assemblies encircle the Golgi apparatus (z-stack intervals: 0.2 µm). 3D tomographic rendering of z-stacked images reveals the spatial relationship between polyQ assemblies and the Golgi stack. PolyQ assemblies encircle the Golgi stack, inducing a localized narrowing of the GM130-stained Golgi ribbon at the encirclement site (yellow arrows: isthmus; blue arrows: polyQ arm; boxed region is magnified in the right panel). **B.** Calibration of GM130-stained Golgi apparatus size in HD and healthy individuals (HD, n = 3; Healthy, n = 3; Student’s t-test, **p < 0.01) **C.** Representative images of clathrin and 3B5H10 antibody immunostaining in the fibroblasts of an HD patient and healthy fibroblasts. The left is a large polyQ assembly (the upper two panels). The inner inserts on the left show angled and rendered views of the spatial relationship between the polyQ assembly and clathrin+ puncta. The right shows the interaction pattern of polyQ with the nucleus and the attachment patterns of clathrin to polyQ assemblies that interact with the nucleus (inner inserts, rendered view [lower] and the conjugation pattern of clathrin with polyQ assembly [upper]). **D.** Lateral views of clathrin, 3B5H10, and phalloidin co-immunostaining reveal that polyQ-conjugated clathrin vesicles reside on the opposite side of F-actin. **E.** Quantification of clathrin⁺ vesicles overlapping with polyQ assemblies, measured via object-object statistics in IMARIS (HD, n = 3; Healthy, n = 3; data represent mean ± SD; Student’s *t*-test, \**p* < 0.05). **F.** Coimmunostaining of 3B5H10 antibody with HAP40 antibody revealed that polyQ assemblies were attached by HAP40. The left panels are sectional views, and the right panels are 3D renderings of the middle panel using the IMARIS surface function. **G.** The count of HAP40 puncta attached to polyQ assemblies in healthy and HD fibroblasts. The distances between HAP40 puncta and polyQ assemblies were measured by IMARIS. HD, n=2; Healthy, n=2. Student’s t-test; Data, mean ± SD. Student t-test; ns, p>0.05 **H.** The 3B5H10 and GM130 antibody immunostaining images in the mitotic fibroblasts, including prometaphase, anaphase, and telophase, were obtained by SIM microscopy. The inserted DAPI images (gray) showed the nuclear chromatin structure. The right panels are a sectional view of the boxed region (anaphase and telophase). **I.** ARF1 and 3B5H10 antibody immunostaining in the fibroblasts revealed that ARF1 preferentially interacts with polyQ assemblies. **J.** The measurement of polyQ assembly-conjugated ARF1 puncta in healthy and HD fibroblasts. Student t-test; \*\**p*<0.001 **K.** GM130 and 3B5H10 immunostaining in BFA-treated fibroblasts reveals that most fragmented Golgi elements dissociate from polyQ. The boxed region in the upper panel is magnified; the lower inset shows a 3D tomographic rendering. **L.** The volume of polyQ assemblies/puncta in fibroblasts after 2 hours and 7 hours of treatment with BFA. The measurements were performed using IMARIS software. (Images, n>6. Sample size: n=2 in each group. One-way ANOVA, \*\**p*<0.01). **M.** The count of 3B5H10-stained assemblies/puncta in the fibroblasts (Images, n=7-11. Sample size: n=2 in each group. One-way ANOVA, \*\**p*<0.01). **N.** Image sequences of Golgi labeled by GOLGI ID Green assay kit in the living fibroblasts of healthy siblings and HD patients (CAG 59, 49, and CAG 49) after glucose starvation (The displays are the inverted channel of the original image). **O.** Golgi fragmentation ratio under glucose starvation across multiple time points in healthy and HD fibroblasts. Fragmentation ratio = (number of Golgi structures per cell at a given time point) / (number of Golgi structures at 0 min) (Two-way ANOVA, \*\**p* < 0.01; images analyzed per cell line, n > 5).

The Golgi apparatus fragments during cell mitosis and reassembles after mitosis (Wei and Seemann, 2009). PolyQ assemblies and the Golgi stack/ribbon fragment during prometaphase (28 cells from 6 samples) and anaphase (23 cells from 6 samples), and reassemble in late telophase (22 cells from 6 samples), and the fragmented Golgi in the mitotic cells conjugates with polyQ **(Fig. 4H).** Consistent with our previous reports (Liu et al., 2024), the polyQ assemblies in HD fibroblasts attached fewer ARF1, and Brefeldin A (BFA) treatments decouple polyQ assemblies from the Golgi apparatus in all fibroblasts **(Fig. 4I-M)**. ARF1, which conjugates to the Golgi after converting its GDP to GTP in a GTP-ATP-dependent manner (D’Souza-Schorey and Chavrier, 2006), couples polyQ assembly to the Golgi (Liu et al., 2024). Thus, it is rational to hypothesize that polyQ assembly in HD fibroblasts is sensitive to energy deprivation. To test this hypothesis, we performed immunostaining on fibroblasts from HD patients and their healthy siblings following 48 h of glucose deprivation, utilizing antibodies against 3B5H10 and GM130. Consistent with the stress-induced alterations in polyQ assemblies observed in both HD and healthy cells, the Golgi ribbons of HD fibroblasts underwent loosening and fragmentation under these conditions. Furthermore, most polyQ fragments and patches were found associated with these fragmented Golgi structures **(Fig. S2A)**. In contrast, healthy fibroblasts maintained a stable Golgi ribbon structure and exhibited fewer small Golgi fragments after 48 h of culture in low-glucose medium **(Fig. S2B)**. Notably, HD patient cells with higher CAG repeat counts (55) displayed more severe Golgi fragmentation than those with lower counts (49) following the 48 h starvation period **(Fig. S2C)**. Additionally, the overall size of the Golgi apparatus in HD fibroblasts decreased more rapidly than that in healthy sibling controls under glucose deprivation **(Fig. S2D)**.

To see *in vivo* Golgi dynamics in a living fibroblast, we continuously monitored fibroblasts (CAG: 49, 55, 19, and 19) stained with a Golgi apparatus-selective dye after glucose starvation or H_2_O_2_ stimulation. The Golgi apparatus in HD patient fibroblasts is more easily fragmented under glucose starvation than in healthy fibroblasts **(Fig. 4N, O)**, whereas the Golgi apparatus in patient and healthy fibroblasts uniformly fragments under H_2_0_2_ stimulation **(Fig. S2E, F).** Taken together, these findings demonstrate that polyQ assemblies form a complex with the Golgi apparatus, a process that is pathologically altered by the presence of mHTT.

Glycosylation reactions mostly happen in the Golgi apparatus (Stanley, 2011). Wheat germ agglutinin (WGA-Alexa-55) staining intensity, which binds sugar residues N-acetylglucosamine and sialic acid, reflects the levels of Golgi glycosylation. We stained HD patient and healthy sibling fibroblasts with WGA conjugated to Cy3 after 3B5H10 and GM130 antibody immunostaining. In 3D SIM images, the HD fibroblasts showed fewer WGA staining puncta in the GM130-stained Golgi region compared to that in healthy fibroblasts **(Fig. S2G).** The overlapped volume of WGA with Golgi or polyQ in HD patient fibroblasts is higher than that in healthy cells **(Fig. S2H, I)**. These findings imply that the glycosylation activity of HD fibroblasts is deficient.

### Impaired polyQ assembly in HD neurons is associated with deficient Golgi function in cortical and striatal neurons

To evaluate these phenotypes in a human neuronal model, we generated iPSCs from the family fibroblasts and differentiated them into 2D GABAergic projection neurons. By day 90 of culture, elongated polyQ assemblies were evident within the projections of striatal neurons from an HD patient (59 CAG repeats), while fragmented polyQ assemblies had deposited within the neuronal nuclei **(Fig. 5A)**. By approximately day 100, intranuclear polyQ assemblies almost completely occupied the nuclei of a subset of striatal neurons across all patient lines (47–59 CAG repeats); in contrast, no nuclear polyQ accumulation was observed in healthy control neurons even after 90 days of culture **(Fig. 5A, B)**. Differential interference contrast (DIC) imaging revealed an intact, distinct, and deeply invaginated cleft where polyQ assemblies resided or traversed **(Fig. S3A)**. Furthermore, 3D tomographic rendering confirmed that these polyQ spindles formed a pronounced groove on the nuclear surface **(Fig. S3A),** underscoring the structural integrity and rigidity of polyQ assemblies. Notably, striatal neurons exhibiting nuclear polyQ accumulation displayed pale-stained nuclei, nuclear membranes distorted or pocked by polyQ, and nuclear fragmentation **(Fig. S3B–F)**. While polyQ fragmentation was also detected in human fetal and pediatric brain samples, it was absent in fresh human brain organoids with no obvious necrosis **(Fig. S4A–C)**. Despite this variation in fragmentation status, the core structural morphology of polyQ assemblies and their specific interaction patterns with the nucleus in brain tissues were identical to those observed in human brain organoids **(Fig. S4B; Fig. 5A, B).** Of note, necrosis-dependent fragmentation of polyQ assemblies was frequently observed in the brain organoids **(Fig. S4D).** Finally, proteomic analysis revealed that key nuclear processes in HD striatal organoids (hStrO), including chromatin and DNA complex assembly, protein–DNA complex formation, DNA packaging, and nucleosome binding, were significantly impaired after 60 days of culture **(Fig. S5A, B)**.

**Fig. 5.**
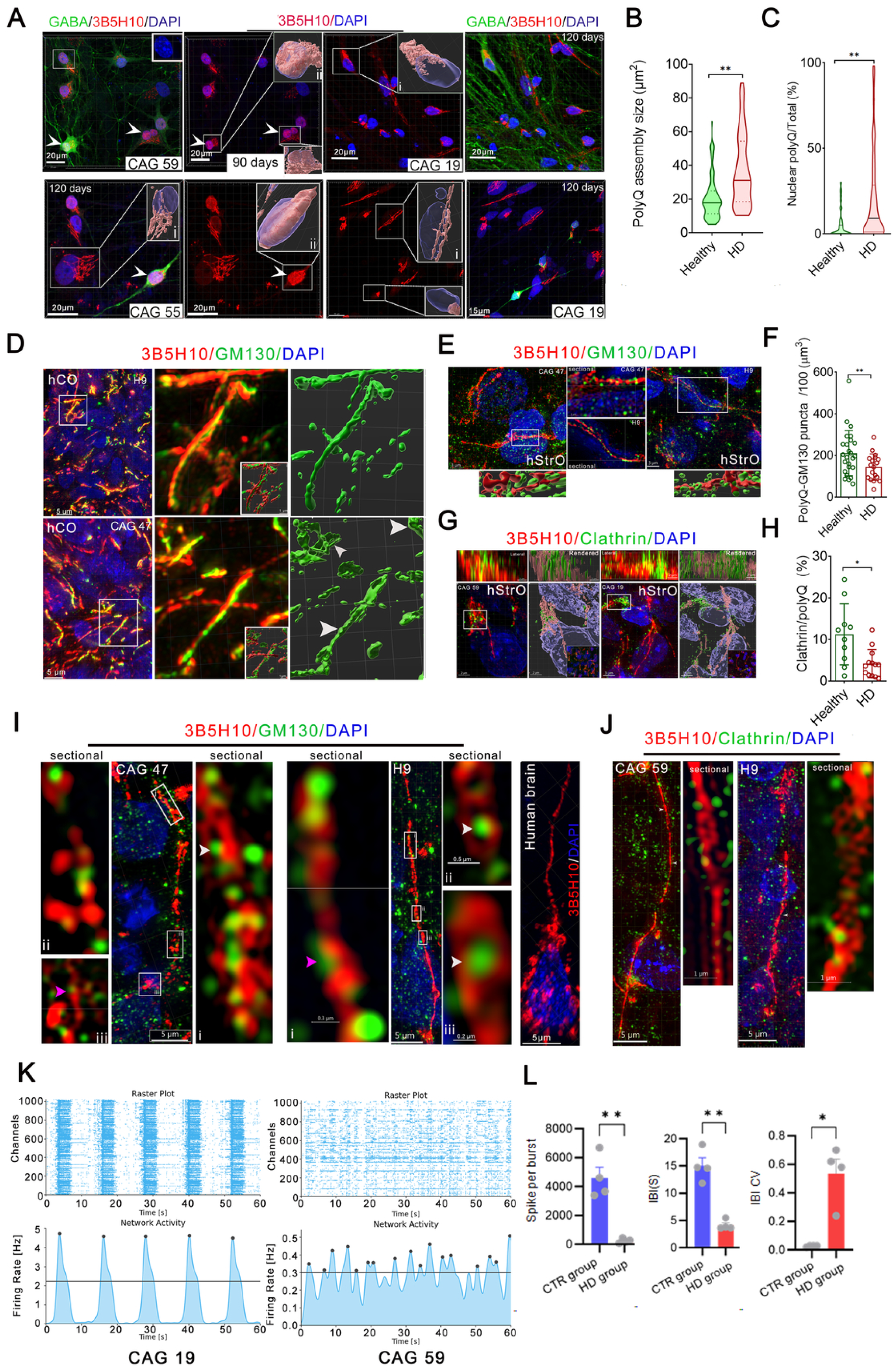
The characteristics of polyQ assembly in the induced striatal and cortical neurons. **A.** The characteristics of 3B5H10-stained polyQ assembly in HD and healthy iPS-derived striatal GABAergic neurons at day 90 and 120 (i, the nucleus without nuclear accumulation; ii, the nucleus with nuclear polyQ deposition). The inner inserts show the 3D tomographic rendering, which revealed the spatial relationship between polyQ and the nucleus (white arrows, the neurons with nuclear deposition of polyQ). **B.** The size of polyQ assemblies in iPSC-derived GABAergic neurons (HD, n=4 [CAG 59, 55, 49, and 47]; Healthy, n=3 [CAG 19, CAG 19, and H9]). Data, mean ± SD. t-test; \*\**p*<0.01. **C.** The percentage of nuclear polyQ volume/total polyQ volume in iPS-derived GABAergic neuronal nuclei cultured over 90 days (HD, n=4 [CAG 59, 55, 49, and 47]; Healthy, n=3 [CAG 19, CAG 19, and H9]). *t*-test; \*\**p*<0.01. **D.** Representative images of polyQ assemblies and Golgi stained by 3B5H10 and GM130 antibodies in hCO (Taken by SIM microscopy; Z-stack interval, 0.2µm). The right panels show the rendered tomography of the Golgi in Imaris 9.8 surface (white arrows indicate the Golgi with an irregular surface). **E.** Representative images of polyQ assemblies and Golgi stained by 3B5H10 and GM130 antibodies in hStrO (taken by SIM microscopy; Z-stack interval, 0.2 µm). The middle panels are a sectional view of the rectangle. The lower panels on the right and left are 3D rendering tomography in IMARIS surface. **F.** Comparing the overlapped volume of the Golgi with polyQ assemblies in HD striatal neurons with that of healthy striatal neurons. The object-object statistic function in Imaris 9.8 measures the overlapped volume (Student’s *t*-test; ** p < 0.01). HD hStrO: CAG 55 and CAG 47; Healthy hStrO: H9, CAG 19. Student’s *t*-test, \**p*<0.05). **G.** The images of polyQ assemblies stained by 3B5H10 and clathrin-coated vesicles stained by clathrin antibodies in hStrOs were taken by SIM microscopy (The upper panel, the lateral view of the boxed region; the inner insert in the rendered image, a larger view of the original image). The renderings are done using the IMARIS surface function. **H.** Comparing clathrin+ vesicle volume overlapped with 3B5H10-stained polyQ assemblies of striatal neurons in the healthy sibling and the HD patient. *t*-test, *, p<0.05. HD hStrO: CAG 55 and CAG 47; Healthy hStrO: H9, CAG 19. Student *t*-test; \**p*<0.05. **I.** 3B5H10 and GM130 antibody immunostaining images and their sectional views in the neurons of hStrO of HD patients and healthy siblings revealed the spatial relationships between polyQ assemblies and Golgi (white arrows, the Golgi in the inner hole of the ring; pink arrows, the scaffolding Golgi; the boxed region, the magnified region in the right or left panel). The right panel is a 3B5H10-stained assembly in a human striatal neuron with a long projection. **J.** 3B5H10 and clathrin antibody staining images of the neurons in hStrO of HD patients and healthy siblings revealed the spatial relationships between polyQ assemblies and clathrin-stained vesicles. The left panel is a sectional view between the two arrows (hStrO at day 80; the two white arrows indicate the segment in the right panel) **K.** The raster plots and network activity of the cortical neurons in HD and healthy hCOs. **L.** Quantification of network features at the burst level in HD and healthy hCOs, including the number of spikes per burst, the interburst intervals (IBI), and IBI CV (CTR group: n=3 hCOs; HD group: n=3 hCOs; mean ± SD, Welch’s t-test, * *p*< 0.05; \*\**p*< 0.01).

We next characterized the structural and spatial relationship between polyQ assemblies and the Golgi apparatus in neurons within human striatal organoids (hStrOs) and human cortical organoids (hCOs) **(Fig. S5A, C)**. In hCOs, polyQ assemblies encapsulated flattened Golgi stacks, and the surface morphology of the Golgi in HD neurons appeared highly irregular **(Fig. 5D)**. Similarly, multiple Golgi vesicles resided within the polyQ assemblies of striatal neurons in hStrOs, which traversed the nuclear surface where a deep gorge exist **(Fig. 5E)**. Quantitative analysis revealed that polyQ assemblies in HD striatal neurons associated with fewer Golgi structures than those in healthy control striatal neurons within hStrOs **(Fig. 5F)**. Conversely, Clathrin^+^ vesicles preferentially distributed along or docked with polyQ assemblies in both HD and healthy control neurons **(Fig. 5G)**. Consistent with our findings in fibroblasts, polyQ assemblies in HD hStrO neurons recruited significantly fewer Clathrin^+^ vesicles and ARF1^+^ puncta than their healthy counterparts **(Fig. 5G; Fig. S6A–C).** Furthermore, the Golgi apparatus in HD neurons displayed a marked reduction in ARF1^+^ puncta localization compared to healthy control Golgi **(Fig. S6D–F)**. In alignment with these localization deficits, ARF1 transcript levels in HD hStrOs were uniformly lower than those in healthy controls at both 60 and 80 days of culture **(Fig. S6G, H)**. Given that striatal projection neurons possess exceptionally long dendrites and axons, we examined extended polyQ assemblies within HD striatal neuronal projections in hStrOs. These structures closely resemble the polyQ assemblies observed in the human striatum and interacted with fragmented Golgi vesicles **(Fig. 5I)**. Specifically, small Golgi structures either resided within the central lumens of ring-like polyQ assemblies or attached to the outer spines of elongated assemblies in hStrO neurons **(Fig. 5I).** Finally, Clathrin^+^ vesicles actively interacted with polyQ assemblies, segregating predominantly along both lateral sides of the assemblies in hStrO neurons **(Fig. 5J)**.

To evaluate the neurophysiological impact of compromised polyQ assembly–Golgi complexes on HD neuronal activity, we performed high-density microelectrode array (HD-MEA) recordings on 80-day-old hCOs. Control organoids exhibited robust, highly synchronized network bursts, resembling the stereotyped activity patterns characteristic of the developing human brain **(Fig. 5K).** In contrast, HD hCOs displayed erratic firing patterns **(Fig. 5K).** Quantitative analysis revealed a significant reduction in spikes per burst, along with an altered inter-burst interval (IBI) in HD hCOs **(Fig. 5L)**. Notably, the elevated IBI CV in HD hCOs signifies a disruption in the periodicity and rhythmic regularity of spontaneous network-wide bursting. Collectively, these functional findings demonstrate that pathogenesis mediated by abnormal polyQ assemblies disrupts the emergence of functional network oscillations and coordinated neuronal firing within cortical circuits.

### Impaired Golgi function in HD striatal neurons

We performed single-cell RNA sequencing (scRNA-seq) analyses of human striatal organoids (hStrOs) derived from hESC-H9, hiPSC-8-12 (CAG 19), and HD hiPSC-7-2 (CAG 59) after 110 days of culture. Transcriptomic data from 38,716 single cells were obtained after quality control and filtering **(Fig. 6A)**. Uniform manifold approximation and projection (UMAP) clustering identified four clusters in hStrOs that expressed cell-type-specific markers, including progenitor (VIM, NES, HES1), neuron (SYT1, DCX, STMN2), astrocytes (S100β, GFAP), and oligodendrocytes (OLIG2, SOX10) **(Fig. 6B, C, and S7A)**. We further plotted the expression of brain region-specific markers in these clusters and found that both control and HD hStrOs are primarily composed of LGE progenitors and GABAergic neurons rather than cortical progenitors and glutaminergic neurons **(Fig. S6C, D)**, consistent with our protocols for generating hStrOs (Chen et al., 2022). Based on the characteristics of Golgi in HD fibroblasts, we compared the transcriptomes of mature GABAergic neurons in the control and HD groups after sorting the GABAergic neuron population using GAD1 and GAD2, specific markers for mature GABAergic neurons **(Fig. S7C-F)**. In the GABAergic neurons, the DEGs of HD hStrO extensively overlapped with those of hiPSC-derived hStrO (CAG 19) and hESC-derived hStrO, both for up-regulated (321 genes) and down-regulated genes (550 genes) **(Fig. 6D)**. Up-regulated genes in GO terms are associated with nuclear activities, including ribonucleoprotein complex assembly, translational initiation, and regulation of translation, which might be related to the nuclear poking of polyQ assemblies or deposition of polyQ into the nucleus.

**Fig. 6.**
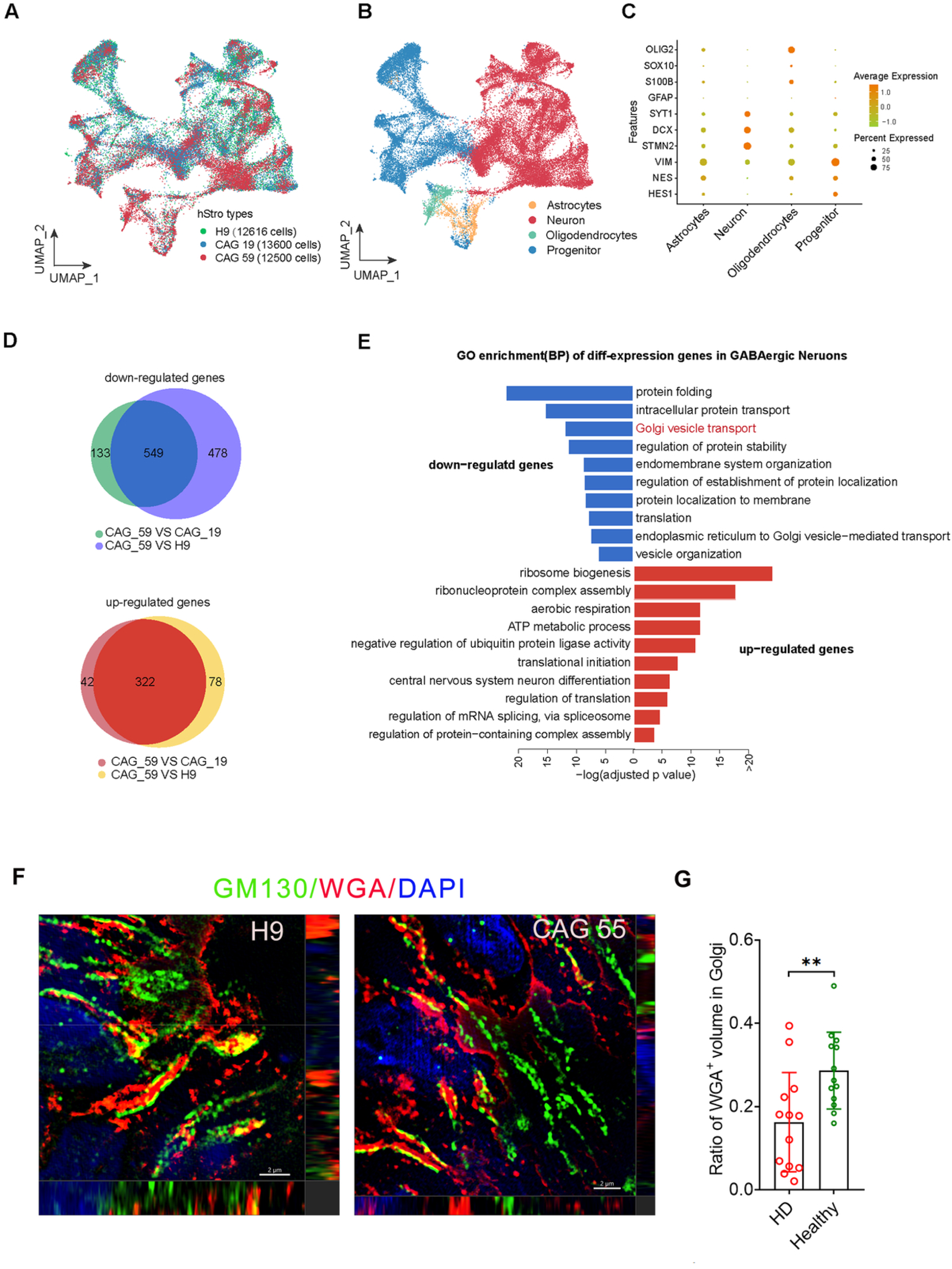
GABAergic neurons from HD patients reduced Golgi- or vesicle-related activities and intensified nucleus-related activities. **A, B**. Uniform manifold approximation and projection (UMAP) plot of all cells colored by their sample origin (A) and by cell types (B). **C.** Dot plot displaying expression of example marker genes. **D.** Venn diagram summarizing the number of overlapping genes differentially expressed in CAG59 vs. CAG19/H9 GABAergic neurons. **E.** GO enrichment analysis of down-regulated (top) and up-regulated (bottom) genes in CAG59 vs. CAG19/H9 GABAergic neurons. **F.** Representative images of WGA conjugated to Cy3 staining in the HD and healthy hStrOs slices immunostained by GM130 antibody. **G.** The ratio of overlapped volume of WGA-stained regions with GM130 antibody-stained regions in hStrOs. Object-object statistics in IMARIS are used to measure overlapped volume. The ratio of overlapped volume=WGA overlapped GM130 antibody-stained regions/GM130-stained regional volume. (n, HD=2 [CAG 55 and 59]; healthy=2; t-test, **p<0.001).

Interestingly, the down-regulated genes in GABAergic neurons of HD hStrOs were notably enriched for GO terms associated with Golgi or Golgi-related functions, including Golgi vesicle transport, endoplasmic reticulum to Golgi vesicle−mediated transport, vesicle organization, endomembrane system organization, intracellular protein transport, protein localization to membrane translation **(Fig. 6E)**. The down-regulated genes in HD hStrOs belong to Golgi matrix proteins, including *GOLGA8A*, a structurally similar protein of GM130 enriched in neurons, *GOLGA4*, and Golgi matrix-associated GTPs that include *RAB6, ARL1/4C*, and *ARF1/3/4.* We also stained the slices of hStrOs with WGA conjugated to Alexa 555 after GM130 antibody immunostaining. Consistent with the WGA staining patterns in fibroblasts, the Golgi region of HD hStrOs showed fewer WGA-labeled puncta compared to that of healthy hStrOs in 3D SIM images **(Fig. 6F)**. The measurement of Golgi-conjugated WGA revealed that the overlapped volume of WGA with GM130-stained Golgi in HD hStrOs was lower than that in healthy hStrOs **(Fig. 6G).** Collectively, these findings showed a systemic disruption of Golgi function in HD striatal neurons.

### Onjisaponin F did not mitigate polyQ nuclear accumulation, nor did ASO treatment normalize HD neuronal firing rates

We selected iPSCs from two HD patients (CAG 59 and 49), one healthy sibling, and the H9 cell line, and differentiated them into striatal neurons in 2D to test the effects of onjisaponin F on striatal neurons. We exposed iPSC-derived GABAergic neurons to a single dose of 20 μM onjisaponin F for 24 h on days 42 and 60. After culturing for another 48 h, we fixed the striatal neurons, assessed polyQ changes, and found that onjisaponin F treatment on days 42 and 60 reduced polyQ assembly volume in HD striatal neurons but not in healthy neurons **(Fig. 7A-C)**. We treated induced HD and healthy striatal neurons with 3 doses of onjisaponin F for 24 h on days 50, 60, and 70. The immunostaining at day 90 showed that 3 doses of onjisaponin (10 µM) decreased the cytoplasmic and total volume of polyQ assemblies in HD striatal neurons but did not reduce the nuclear deposition of polyQ; the immunostaining at day 110 revealed that 3 doses of onjisaponin (20 µM) did not change the volume of polyQ assembly in healthy striatal neurons **(Fig.7D-E).**

**Figure 7.**
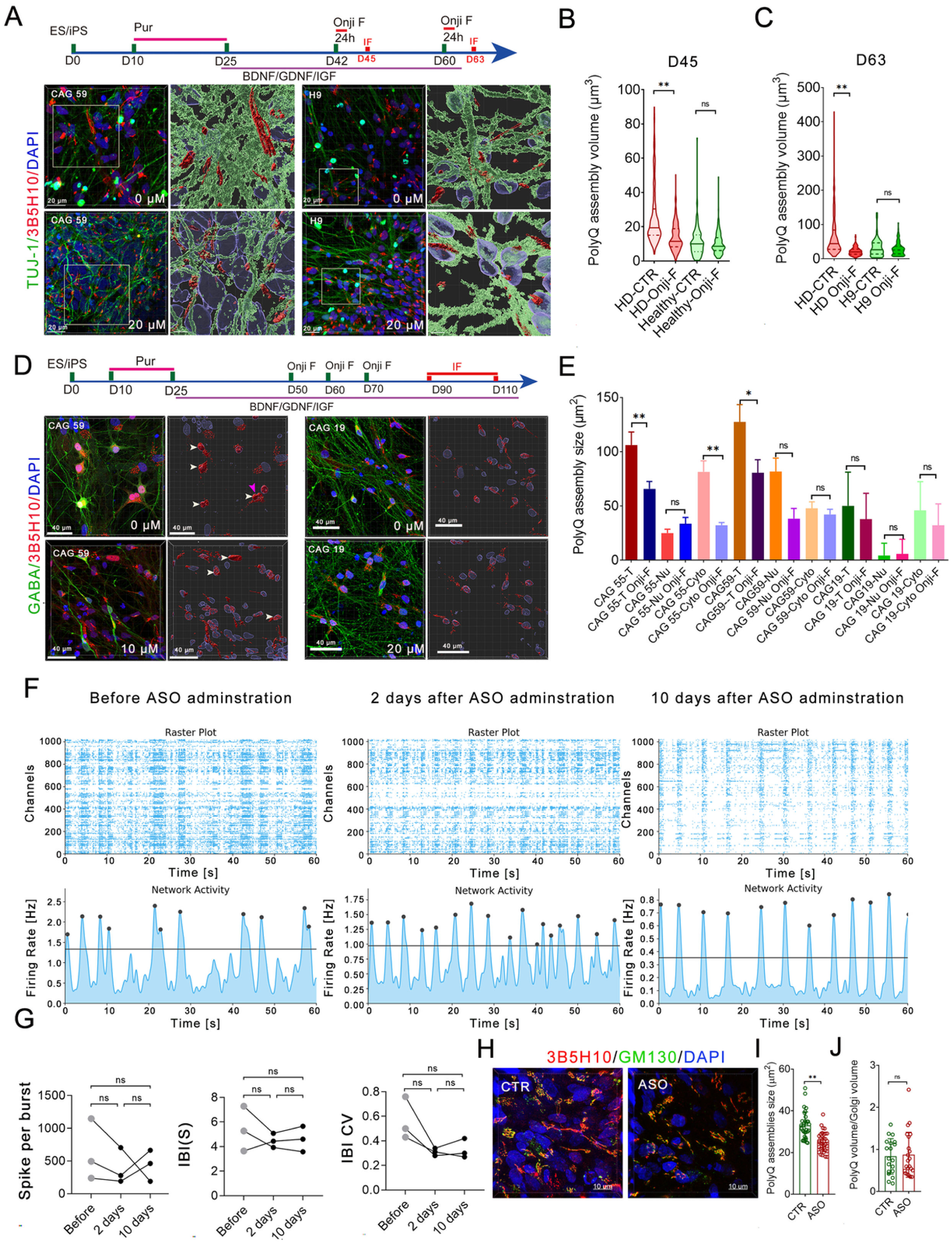
Onjisaponin F and ASO treatment reduce polyQ assembly but do not mitigate polyQ nuclear accumulation and aberrant neuronal firings. **A.** The schematics of treatment (upper) and the representative images of polyQ assemblies stained by 3B5H10 antibodies in HD and healthy striatal neurons at 63 days, treated with one dose of 20 µM onjisaponin F for 72 h. **B, C**. Changes in polyQ assemblies in HD and healthy striatal neurons at days 45 and 63, treated with onjisaponin F for 72 h (HD, n=3 [CAG 59, 55, and 49]; Healthy, n=2 [CAG 19, CAG 19]; One-way ANOVA; ** <0.01). Data, Mean ± SD. **D.** The schematics of treatment (upper) and the representative images of polyQ assemblies stained by 3B5H10 antibodies in HD and healthy striatal neurons treated with 3 dosages of 10 µM onjisaponin F treatment at day 50, day 60, and day 70 (CAG 59). Neurons were fixed at day 90−98 and stained with a 3B5H10 antibody (white arrows, nuclear polyQ deposition). 3 doses of 20 µM onjisaponin F treatment at days 50, 60, and 70 did not alter the polyQ assembly in healthy neurons (CAG 19). The healthy neurons were fixed at 110 days and immunostained with 3B5H10 and GABA antibody. **E.** Calibrating the total (T), cytoplasmic (Cyto), and nuclear (Nu) polyQ assembly volume in HD and healthy striatal neurons treated with 3 dosages of onjisaponin and cultured for 90 or 110 days (One-way ANOVA; ** *p*<0.01. Data, means± SD). **F.** The raster plots and network activity in the cortical neurons of Day 80 hCOs before and after ASO treatments. **G.** Quantification of network features at the burst level in the cortical neurons of hCOs, including the number of spikes per burst, the interburst intervals (IBI) [s], and IBI CV (Friedman test, ns, *p* > 0.05; hCOs, n=3) **H.** The immunostaining of ASO-treated and control (CTR) HD hCOs with 3B5H10 and GM130 antibodies. Super-resolution images were taken by SIM. **I.** Measuring the size of polyQ assembly in ASO-treated and control (CTR) hCOs (HD: n=2 [CAG 49 and 57]; hCOs, n=4, t-test, \*\**p*<0.001). **J.** Comparing the polyQ volume/Golgi volume ratio in ASO-treated and control (CTR) hCOs (HD: n=2 [CAG 49 and 57]; hCOs, n=4; t-test, ns, *p*>0.05).

Given that ASOs have demonstrated efficacy in promoting mHTT degradation across experimental models (Kordasiewicz et al., 2012) and have reduced the polyQ assembly size in human fibroblasts. We utilized ASOs as a benchmark to evaluate their therapeutic impact on the polyQ assembly-driven phenotypes in hCOs. Following treatment, we observed downward trends in both spikes per burst and the IBI **(Fig. 7F, G).** The IBI coefficient of variation (CV) also trended lower, suggesting a subtle shift toward improved firing patterns **(Fig. 7F, G).** However, these alterations failed to reach statistical significance across 10 days **(Fig. 7F, G)**. The analysis of polyQ, Golgi, and ARF1 by immunostaining revealed that ASO treatment reduced the size of polyQ assemblies in HD-hCOs but did not alter the Golgi tomography and did not increase the recruitment of ARF1 and Golgi to polyQ assemblies **(Fig. 7H-J**; **Fig. 8A-D**). These findings demonstrate that, under the tested conditions, the ASO treatment was insufficient to achieve a significant functional recovery of the HD-hCO network dynamics and rescue the Golgipathy in HD-hCOs.

## Discussion

This study redefines huntingtin (HTT) polyQ structures, previously considered strictly pathogenic (Truant et al., 2008), as large, dynamic, and highly organized structural networks that form a coupled polyQ-Golgi complex. Utilizing patient-derived cells from a large HD pedigree and state-of-the-art imaging, we demonstrate that these polyQ assemblies consist of multiple elongated, parallel, and interfused spindles. This architectural network encircles the Golgi apparatus and binds clathrin-coated vesicles, establishing a structurally and dynamically interdependent polyQ assembly-Golgi complex. Crucially, the presence of mHTT with an expanded polyQ tract alters the responsiveness of this network to both energy deprivation and autophagy-enhancing pharmacological agents. Furthermore, mHTT destabilizes the Golgi apparatus by impairing ARF recruitment, ultimately driving a distinct Golgipathy in HD patients.

The mammalian Golgi ribbon comprises up to 100 flattened stacks—each composed of 4–11 aligned cisternae, where newly synthesized lipids and soluble cargo proteins are processed and sorted (Klumperman, 2011; Liu et al., 2021). These stacks are interconnected by tubular bridges and supported by the Golgi matrix, a ribosome-free protein network that maintains the organelle’s structural integrity (Ravichandran et al., 2020). Intriguingly, the spatiotemporal relationships among polyQ assemblies, the Golgi apparatus, clathrin-coated vesicles, and ARF1 suggest that rigid polyQ assemblies function as structural components of the Golgi matrix. In this capacity, they likely provide a scaffold that regulates Golgi morphology, assembly, and sorting activities. ARFIP2 interacts with multiple guanine nucleotide exchange factors (GEFs), including ARF1, ARF3, ARF5, and ARF6 (Wu and Zhou, 2009), and directly binds to HTT (Peters et al., 2002). Thus, the ARFIP2-ARF complex represents a rational candidate for mediating the interaction between polyQ assemblies and the Golgi apparatus. This model is further supported by our finding that BFA decouples the polyQ assembly-Golgi complex, whereas mitotic Golgi fragmentation does not, reinforcing the specific role of the ARFIP2-ARF1 complex in mediating this structural assembly.

Golgipathy, including structural and functional abnormalities, is common in neurodegenerative diseases (El Ghouzzi and Boncompain, 2022; Klumperman, 2011; Liu et al., 2021; Passemard et al., 2017). The correct Golgi positioning is critical for cellular polarization and differentiation (Yadav and Linstedt, 2011). The precise localization of the Golgi apparatus mediates the simple spindle-like cells into dentate granule cells (DGCs) (Rao et al., 2018); GM130 deletion causes neurodegeneration and ataxia (Liu et al., 2017). Pathogenic *GORASP1 and ARF3* variants cause a neurodevelopmental disorder with neurosensory, neuromuscular, and skeletal abnormalities by affecting the cell cycle and glycosylation (Fasano et al., 2022; Lebon et al., 2025; Sakamoto et al., 2021). Previously, we reported that polyQ assemblies in HD cortical neurons altered corticogenesis by impairing ARF1 recruitment to the Golgi apparatus in the neuroepithelium(Liu et al., 2024). The glycosylation deficiency in HD cells indicates that endogenous polyQ expansion impaired Golgi glycosylation, supporting the Golgipathy of HD patients.

Striatal projection neurons (SPNs), which characteristically display a prominent, elongated polyQ assembly alongside nuclear polyQ accumulation, are among the earliest cells lost in HD patients (Walker, 2007b). Given that huntingtin (HTT) operates as a structurally rigid protein (Dougan et al., 2009; Guo et al., 2018; Saudou and Humbert, 2016; Sivaramakrishnan et al., 2008; Tabrizi et al., 2020), maintaining the integrity of these macromolecular structures poses a high energetic demand. Specifically, the recruitment and activation of ARF1 at the Golgi membrane is an ATP-dependent process (D’Souza-Schorey and Chavrier, 2006). While HD is traditionally associated with systemic energy deficits, certain compensatory responses drive elevated glucose metabolism and glycolytic reprogramming under cellular stress (Tang et al., 2013). Grounded in these homeostatic compensation mechanisms, the deficient ARF1 recruitment to the Golgi apparatus seen in HD may act as a driver for hypermetabolic shifts, as the polyQ assembly-Golgi complex in HD striatal neurons requires additional ATP to sustain its structure and function.

Intriguingly, treatment with the potent autophagy enhancer Onjisaponin F preferentially clears polyQ assemblies in both HD fibroblasts and SPNs without altering the underlying configuration of the remaining assemblies. However, this clearance fails to rescue nuclear polyQ deposition, a classical hallmark of HD pathology (DiFiglia et al., 1997). Considering the precise spatial proximity of the polyQ assembly to the neuronal nucleus, alongside the inherently rigid biophysical nature of polyQ, our data strongly suggest that the physical fragility of the HTT assembly, rather than its absolute size, governs nuclear accumulation in long-term cultured striatal neurons. This size-independent paradigm is further supported by therapeutic interventions; a well-characterized antisense oligonucleotide (ASO) successfully reduced the total volume of polyQ assemblies in HD cells, yet failed to correct aberrant neuronal firing in HD cortical networks. Taken together, these findings indicate that the intrinsic structural stability of the polyQ assembly-Golgi complex, rather than the macroscopic size of the assemblies themselves, acts as the primary driver of pathological neuronal activity.

Our study reveals the structure of polyQ aggregates and their spatial-temporal relationship with the Golgi apparatus, and demonstrates that mHTT destabilizes the polyQ assembly-Golgi complexes, dismantles the recruitment of Golgi activity-related proteins, leading to Golgipathy in HD. Thus, improving Golgi function and stabilizing polyQ assembly in HD might represent another approach to treating HD.

### The limitation of this study

While our study establishes a novel structural framework for HTT biology, several limitations warrant consideration. First, the precise stoichiometry of mutant versus wild-type HTT (mHTT-to-wtHTT) likely modulates the functional deficit observed within the polyQ-Golgi complex. However, the exact spatial incorporation pattern of mHTT into these macromolecular polyQ assemblies remains to be elucidated. Second, although we characterized the structural fragility of polyQ assemblies under cellular stress in HD models, it remains unclear how this structural instability drives subsequent nuclear polyQ deposition. Future investigations deploying super-resolution and longitudinal live-cell imaging in HD models will be instrumental to track the real-time assembly, dynamics, and transition of these cytoplasmic complexes into nuclear inclusions. Third, human cells express eight other distinct polyQ-containing proteins—including Atrophin-1, Androgen Receptor, Ataxin-1, -2, -3, -7, TATA box-binding protein, and CaV2.1, whose expansion triggers neurodegenerative diseases (Villavicencio Gonzalez and Zoghbi, 2026). Whether any of these heterologous polyQ-bearing proteins co-aggregate or participate in scaffolding these specific macromolecular assemblies remains unknown.

## Materials and methods

### Ethics

The ethics of this HD work were approved (No. 28) by the Ethics Committee of the Institutes of Biomedical Sciences at Fudan University. All HD family members who participated in this study signed a written consent form translated into Mongolian, and consent to enter this study and authorization for publication were obtained. Dr. Li Li collected the human brain samples. The ethics approval number was obtained from the Huashan Hospital Ethics Committee (KY2018-310.01).

### Human fibroblast and blood sample collection

After obtaining consent, a surgeon collected skin samples from the upper arm using a Skin Punch (diameter: 0.3 x 0.3 cm). After collecting samples, a nurse and doctor visited the patients daily for 7 days to prevent inflammation. The tissues were harvested into a tube with culturing medium supplied with antibiotics. After being transported to a lab, tissues were moved to a Biosafety Cabinet, sliced into smaller fragments with surgical knives, and then attached to dishes with forceps and cultured in a small amount of medium. After fully attaching to dishes, the tissues were placed in sufficient medium and cultured continuously for several weeks to obtain sufficient fibroblasts.

## Author contributions

HS and LM set up this study. HS and LM performed most of the experiments; HS analyzed the data and prepared the manuscripts; XC, WY, YL, BH, and JX performed image analysis and immunostaining. XC, LY, and LM constructed organoids.

## Financial supports

This work was supported by the National Natural Science Foundation of China (grant no. 32370852 and U24A2014 to LM) and the National Key Research and Development Program of China (grant no. 2021YFA1101302 to LM)

## Supplementary Figure legends

**Figure S1.**
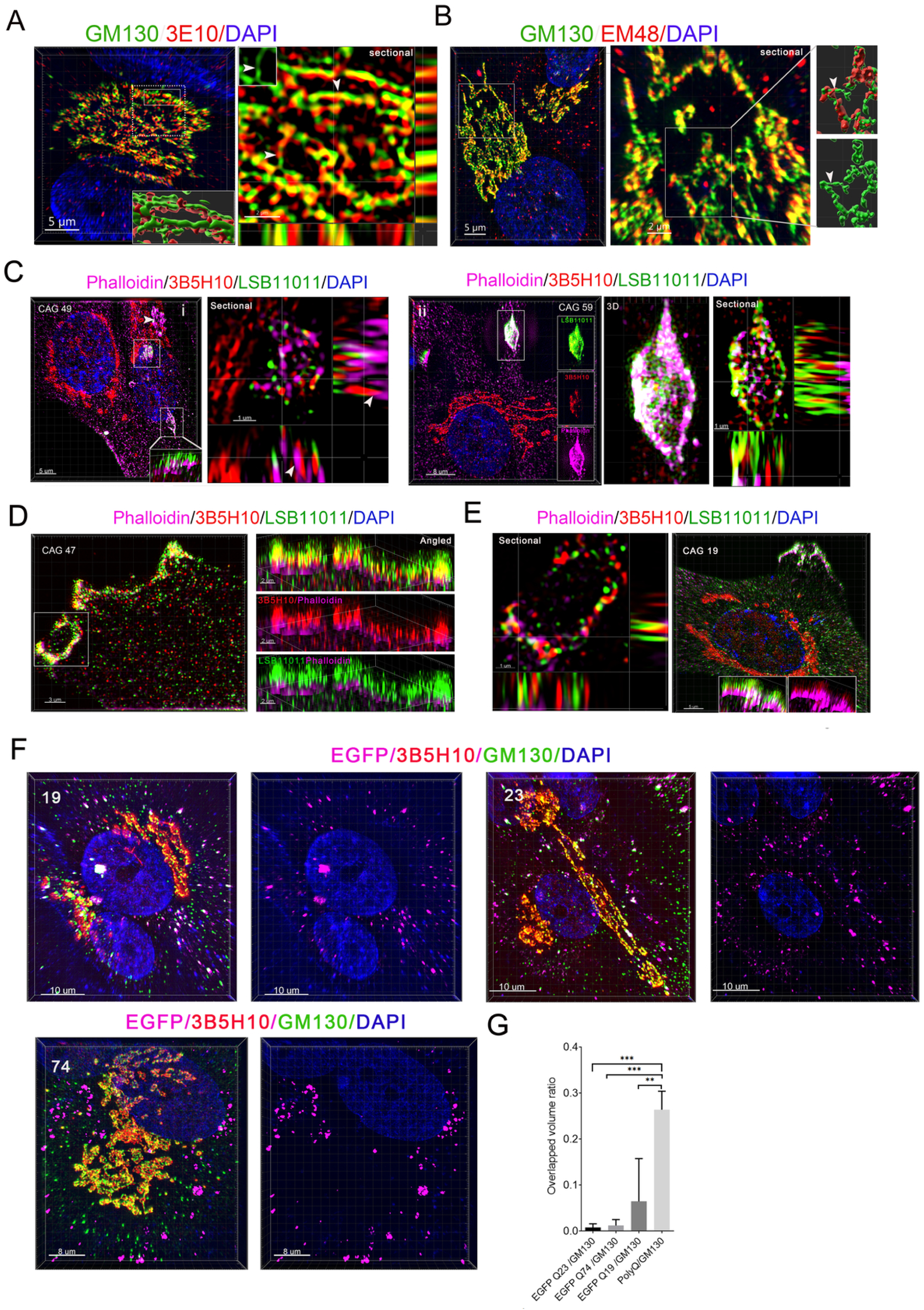
PolyQ assemblies potentially composed of a large amount of HTTs. **A, B.** Coimmunostaining of GM130 antibody with 3E10 or EM48 antibody in the fibroblasts revealed that polyQ assemblies potentially contain a decent amount of HTTs. The right panel is the sectional view of the dashed rectangle (A). The boxed region on the right is the rendering region in the inner insert (right below) (A). The boxed region on the right is the magnified region in the middle panel (B). The boxed region in the middle is the rendering region in the right panel (B). The white arrows indicate the isthmus-like structure of the Golgi (A, B). **C.** LSB11011 (C-terminal of HTT) and 3B5H10 (N-terminal) antibody and phalloidin immunostaining revealed that actin filaments are enriched on the border region of polyQ assembly that uniformly stained by C-terminal antibody and the N-terminal of HTTs docked onto F-actin (i) or a vesicle-like structure of HTTs in the fibroblast present in cells and the N-terminal of HTTs docked onto F-actin (ii) (the boxed region indicated the sectional view in the right panel; white arrows, the interaction point of HTTs with F-actin; the inner panels in ii are the spliced channels; white arrows, the interacting patterns of HTTs with F-actin). **D, E.** LSB11011 and 3B5H10 antibodies, and phalloidin co-immunostaining in the fibroblasts revealed a long assembly of HTTs on the ruffling membrane. The right panel shows a 3D image (angled view) revealing the alignment pattern of HTTs on membrane curvature and the ruffling membrane (D), showing that the N-terminus of HTTs docked onto F-actin. The image of sectional views in the right panel revealed the interaction pattern of HTTs with F-actin filaments on membrane curvature (E). The inner inserts are an angled view of a 3D image (E). **F.** The 3D view images of the fibroblasts transfected with the vector of *HTT* first exon (Q19, Q23, and Q74) were immunostained with 3B5H10 and GM130 antibodies. **G.** The overlapped volume of the first exon fragments of HTT tagged by EGFP (Q19, Q23, and Q74) with the GM130-stained Golgi apparatus.

**Figure S2.**
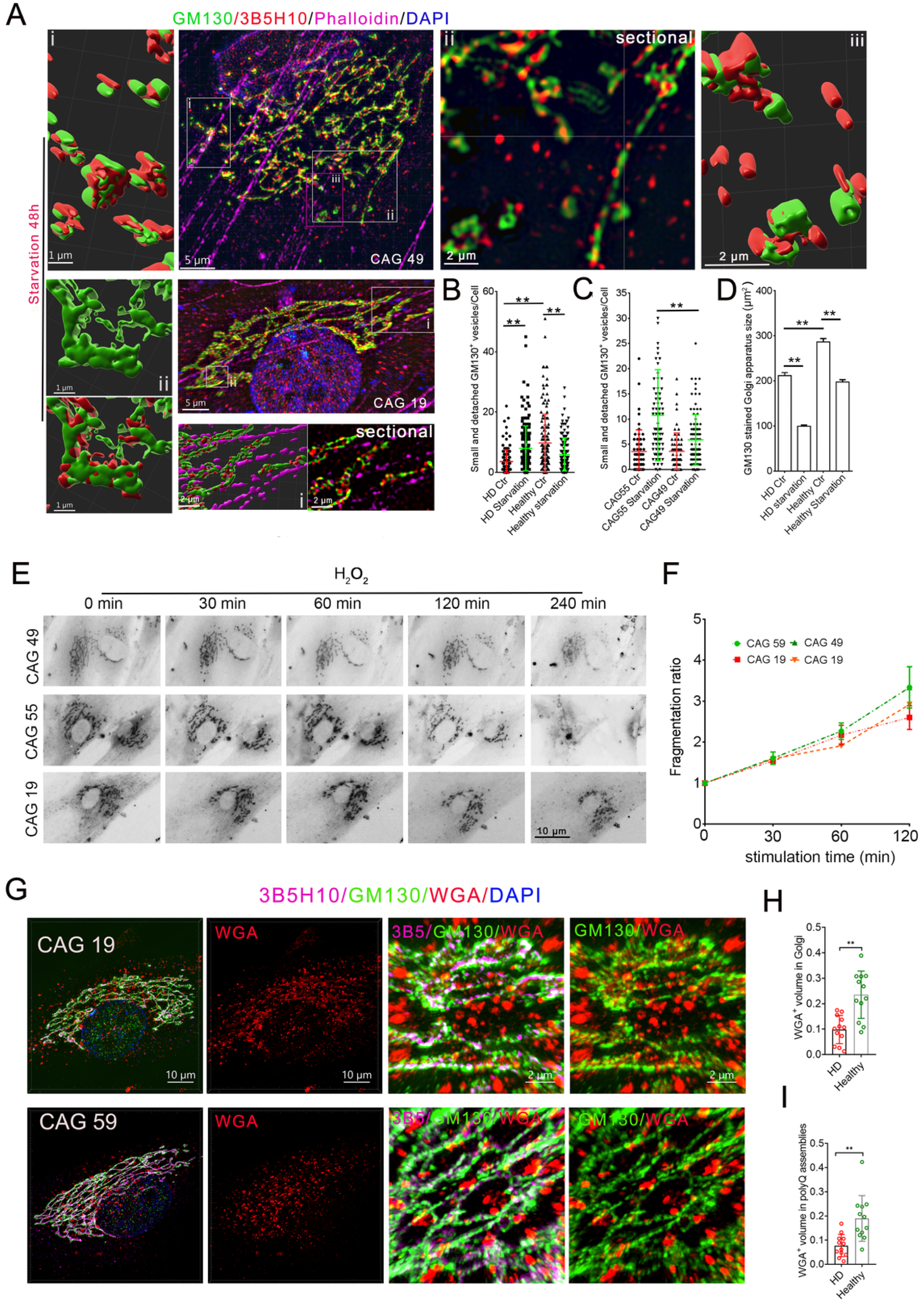
Structural alterations of the Golgi apparatus in HD and healthy control fibroblasts under metabolic and oxidative stress. **A.** Representative structured illumination microscopy (SIM) images of fibroblasts co-stained with 3B5H10 (polyQ) and GM130 (Golgi apparatus) antibodies following 48 h of glucose starvation. Right panels display 3D tomographic reconstructions illustrating the spatial relationship between polyQ assemblies/puncta and the Golgi apparatus. Boxed regions denote magnified views or specific 3D-rendered areas. **B, C.** Measurements of Golgi fragmentation following 48 h of glucose starvation in fibroblasts from healthy siblings (n = 2 independent lines, 19 CAG repeats) and HD patients (n = 2 [49 and 55 CAG repeats]). Two-way ANOVA; **, p < 0.01. Cell count per experimental group: n > 120 cells. **D.** Changes in GM130-stained Golgi sizes after 48 h glucose starvation in fibroblasts from healthy siblings (two, CAG 19) and HD patients (CAG 55 and CAG 49). HD/healthy ctrl: fibroblasts cultured in high-glucose medium; HD/healthy starvation: fibroblasts cultured in low-glucose medium for 48 h. One-way ANOVA; \*\**p*<0.01. The number of images per cell line: n>20. Cell count in each group: n>400. **E.** Representative time-lapse live-cell image sequences tracking Golgi morphology in HD and healthy control fibroblasts at the indicated time points under oxidative stress conditions. **F.** Quantification of the Golgi fragmentation ratio over time under hydrogen peroxide H_2_O_2_ stimulation. The fragmentation ratio was calculated as: (number of discrete Golgi elements per cell at a given time point) / (number of discrete Golgi elements per cell at 0 min). Statistical significance was evaluated using a two-way ANOVA; *, *p* < 0.05; **, *p* < 0.01. **G.** Representative WGA (Alexa 555) staining in the HD and healthy fibroblasts immunostained with 3B5H10 and GM130 antibodies. Images were taken under SIM microscopy with 0.4 µm intervals in Z-stacks. The rectangles in the left panels are the magnified regions in the right two panels. **H, I.** The ratio of overlapped volume of WGA-stained regions with GM130- or 3B5H10-antibody-stained regions (n, HD=2 [CAG 57 and 59]; healthy=2). Object-object statistics in IMARIS are used to measure overlapped volume. The ratio of overlapped volume=WGA overlapped GM130- or 3B5H10 antibody-stained regions/GM130 stained or 3B5H10-stained volume. . Student t-test; \*\**p*<0.01.

**Figure S3.**
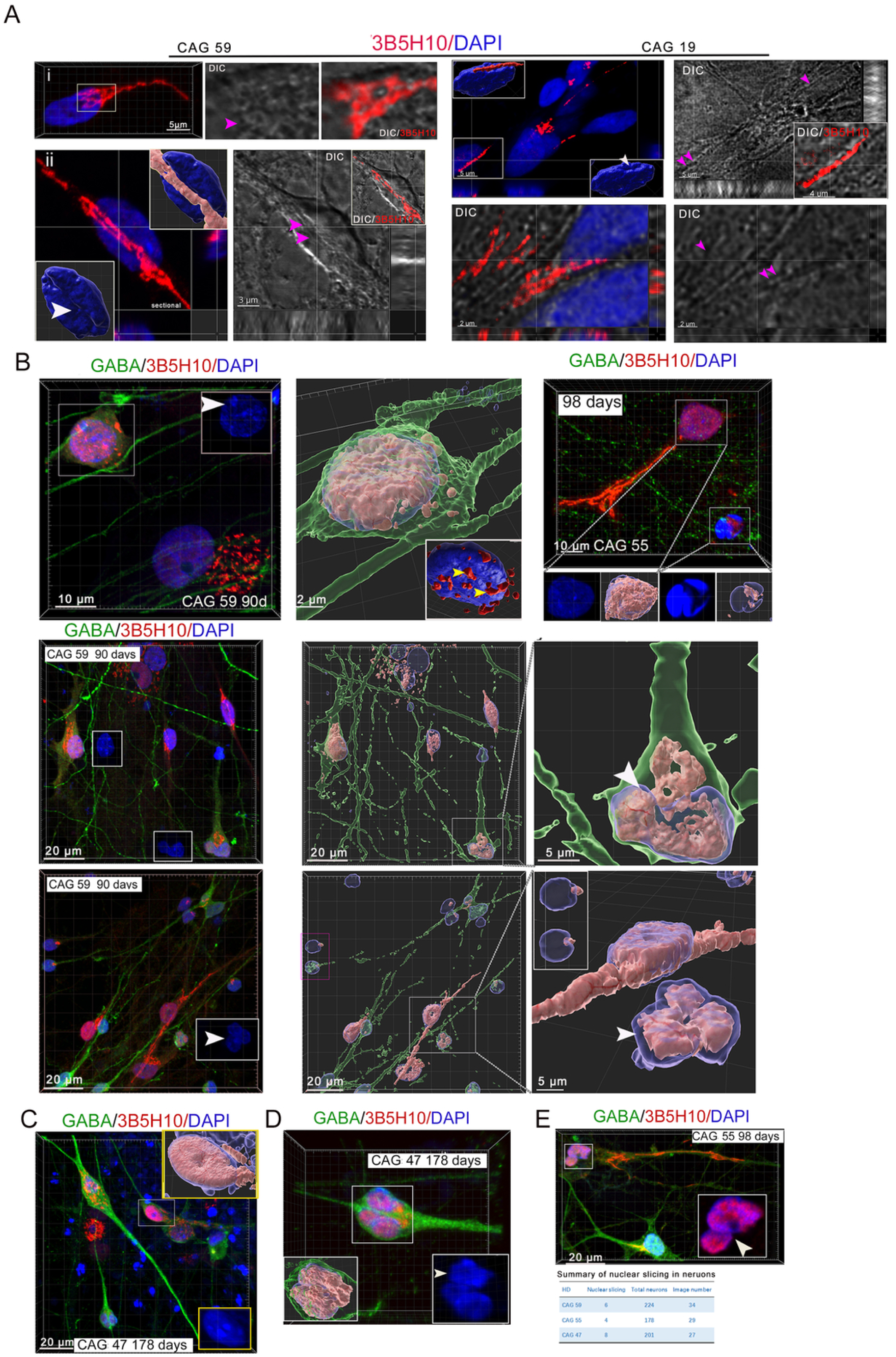
Structural characteristics of polyQ assemblies and intranuclear polyQ deposition in induced neurons. **A.** DIC microscopy and 3D-rendered tomographic reconstructions demonstrating that elongated polyQ assemblies form a deep cleft or groove within the cytoplasm or along the nuclear envelope of induced neurons cultured for 75 days (19 and 59 CAG repeats). Pink arrows (i, ii) denote the deep nuclear grooves/clefts; white arrows indicate nuclear surface indentations visualized via IMARIS surface-rendering software. High-magnification inner inserts and 3D tomography illustrate the spatial relationships. **B.** PolyQ deposits into the nucleus and pokes the nuclear membrane of striatal neurons (white arrow, the morphology of the nucleus; yellow arrows, nucleus poked by polyQ assemblies). The lower left in the right panel is an apoptotic cell (CAG, 55). **C.** PolyQ precipitates occupied the nucleus of GABAergic neurons cultured for 178 days (CAG, 47). **D, E.** HD patient striatal neurons derived from iPSCs cultured for 90 days (2D culture) (CAG 59), 98 days (CAG 55), and 110 days (CAG 47) have a spliced nucleus (white arrows, spliced nucleus). The summary table of nuclear splicing.

**Figure S4.**
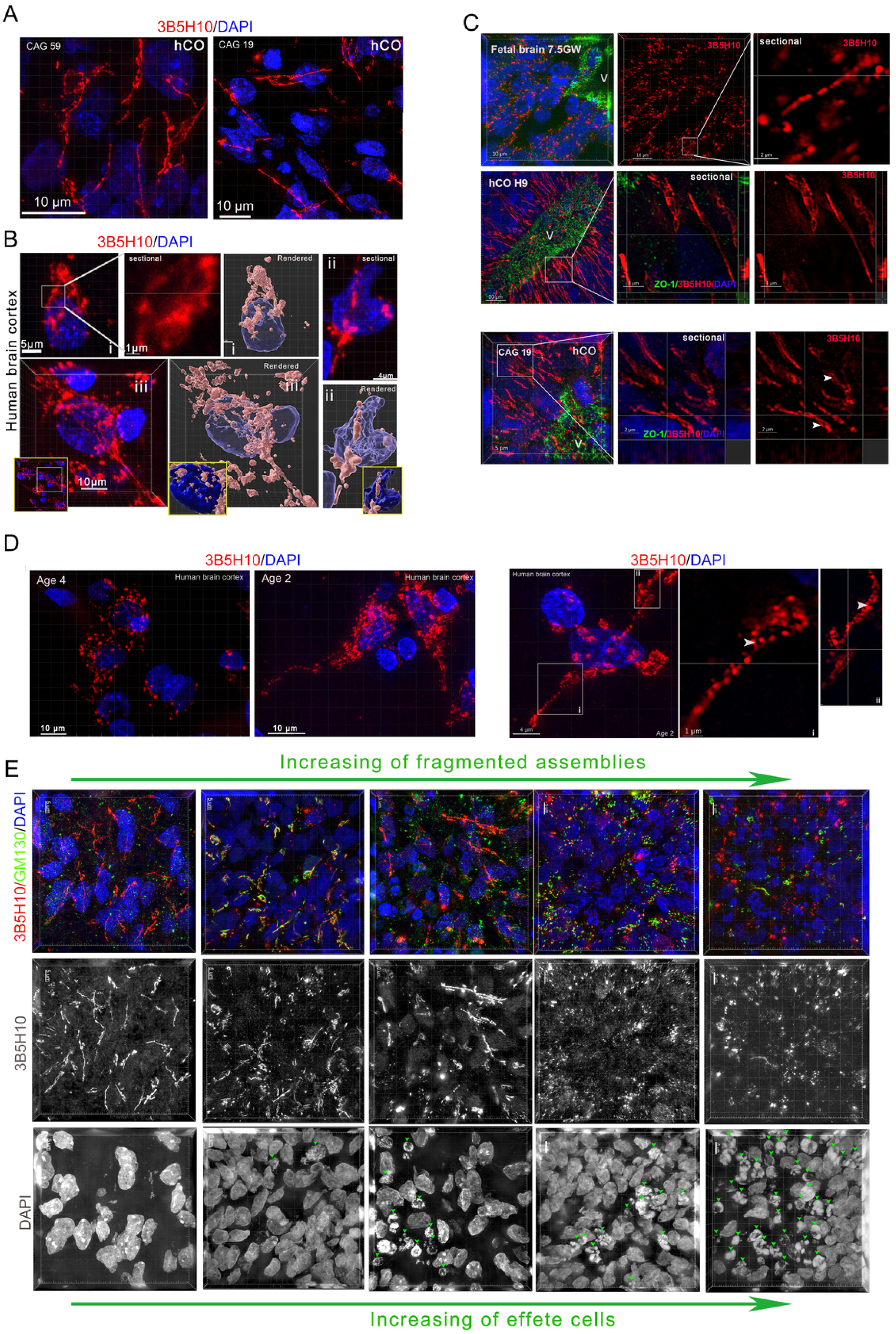
PolyQ assemblies in human fetal and child brains and human brain organoids with necrotic cells. **A.** Representative immunofluorescence images showing the typical elongated, continuous, and structurally intact morphology of polyQ assemblies within the neurons of hStrOs. **B.** PolyQ assemblies stained by 3B5H10 antibody in fresh surgical brain tissues collected from a 4-year-old glioblastoma patient. Three representative appearances of polyQ assemblies in human brain cells. The middle panel (i) is a sectional view that shows a ring-like structure; the left corner insert in image ii is a larger view; the rendered tomography shows the nuclear surface. **C**.3B5H10 and ZO-1 antibodies immunostained intact polyQ assemblies in the pseudostratified neuroepithelium of the neural tube in the human fetal brain at 7.5 gestation weeks, and hCOs from HD patients, H9, and healthy siblings showed that the polyQ assemblies in the neuroepithelial cells are intact and long and formed by a small ring-like unit, and paralleled the apical-basal axis of cells. **D.** Fragmented/discontinuous polyQ assemblies in freshly surged human brain samples from cortex (tissues from 4-year-old and 2-year-old glioblastoma patients). **E.** The fragmentation of polyQ assemblies depends on the necrotic cells in the surrounding regions (green arrows, necrotic cells).

**Figure S5.**
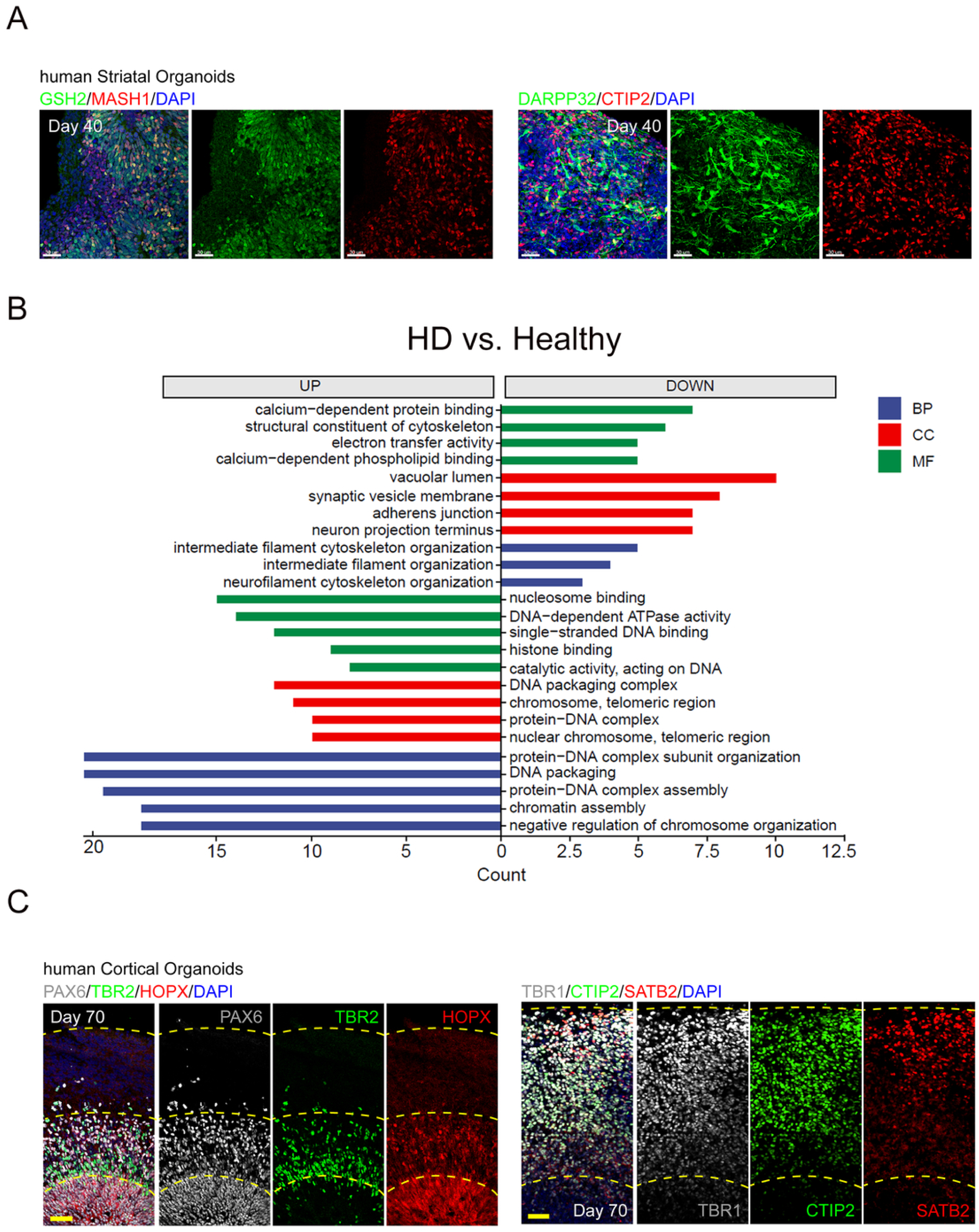
Impaired nuclear activities in HD striatal organoids. **A.** The validation of hCO and hStrOs by immunostaining with GSH2/MASH1 or DARPP321/CTIP2 antibodies. **B.** GO analysis of differentially expressed proteins in HD striatal neurons derived from iPSCs and healthy GABAergic neurons derived from iPSCs at 60 and 100 days (CAG 55 vs. H9 and CAG 19). **C.** The validation of hCO by immunostaining with PAX6/TBR2 /HOPX or TBR1/CTIP2/SATB2 antibodies.

**Figure S6.**
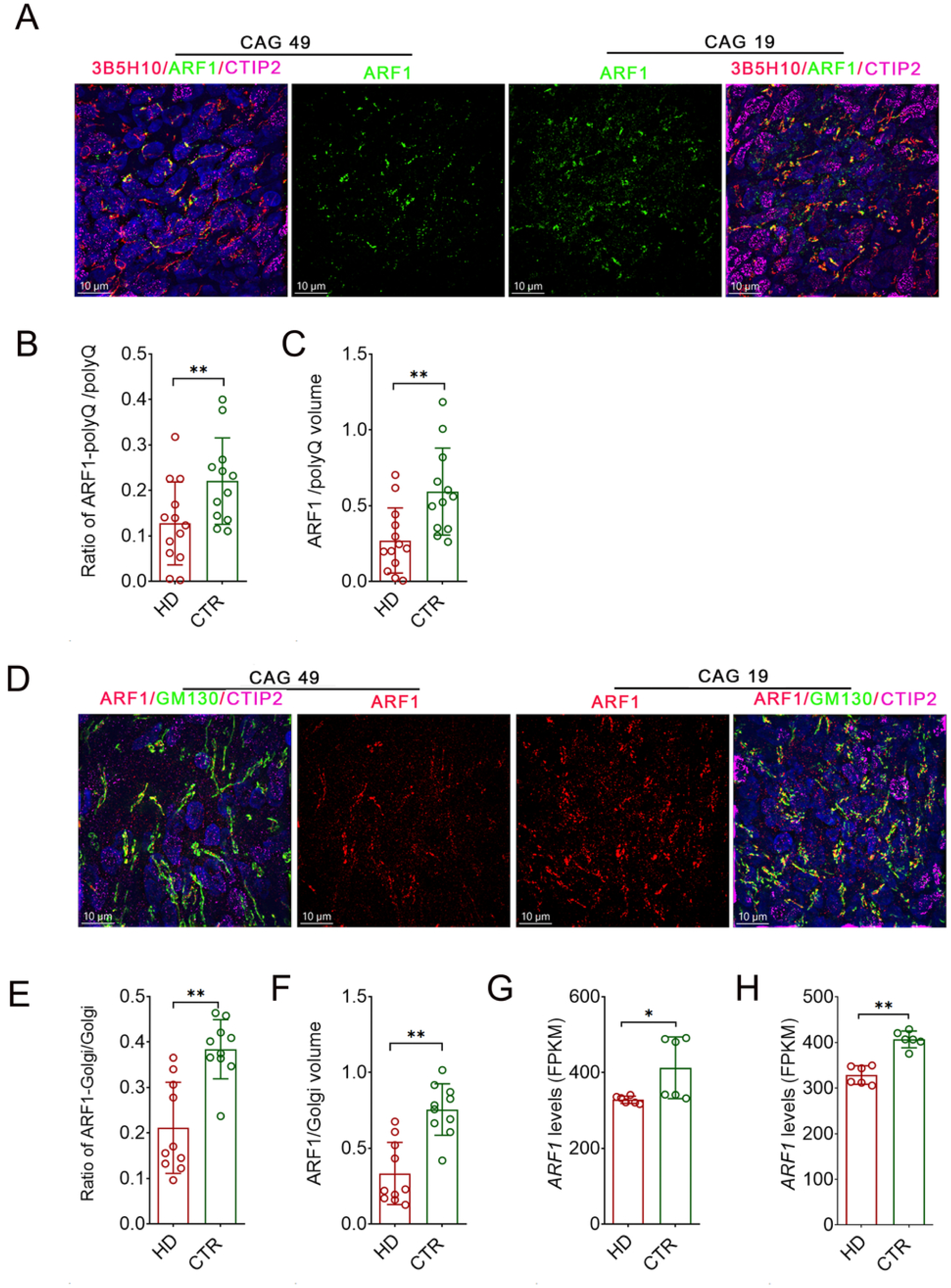
ARF1 levels were reduced in HD neurons. **A.** The immunostaining results of ARF1, CTIP2, and 3B5H10 antibodies in the neurons of hStrOs (CAG 49 and CAG 19). The middle panels showed ARF1 signals. **B, C.** The overlapped volume of ARF1 with polyQ in the HD and healthy neurons in hStrOs (G). The ratio of ARF1 volume/polyQ volume in CTIP2 + regions. The surface function of IMARIS was used to calibrate volume. Object-object statistics in IMARIS measure the overlapped volume. *t*-test, \**p*<0.05; \*\**p*<0.01. **D.** The immunostaining results of ARF1, CTIP2, and GM130 antibodies in the neurons of hStrOs (CAG 49 and CAG 19). The middle panels show ARF1 signals. **E, F.** The overlapped volume of ARF1 with the Golgi in the HD and healthy neurons of hStrOs. The ratio of ARF1 volume/Golgi volume in CTIP2 + regions. The surface function of IMARIS was used to calibrate volume. Object-object statistics in IMARIS measure the overlapped volume. *t*-test, ** *p*<0.01. **G.** Bulk RNA-seq data of HD hStrOs (CAG 55 and 59) and healthy hStrOs (CAG 19 and H9) at 60 days revealed *ARF1* transcription levels*. Student t-test; *p<0.05; **p<0.01.* **H.** The bulk RNA-seq data of HD hStrOs (CAG 55 and 59) and healthy hStrOs (CAG 19 and H9) at 80 days revealed *ARF1.* Student *t*-test; *\*\*p<0.01.* CTR, healthy.

**Figure S7.**
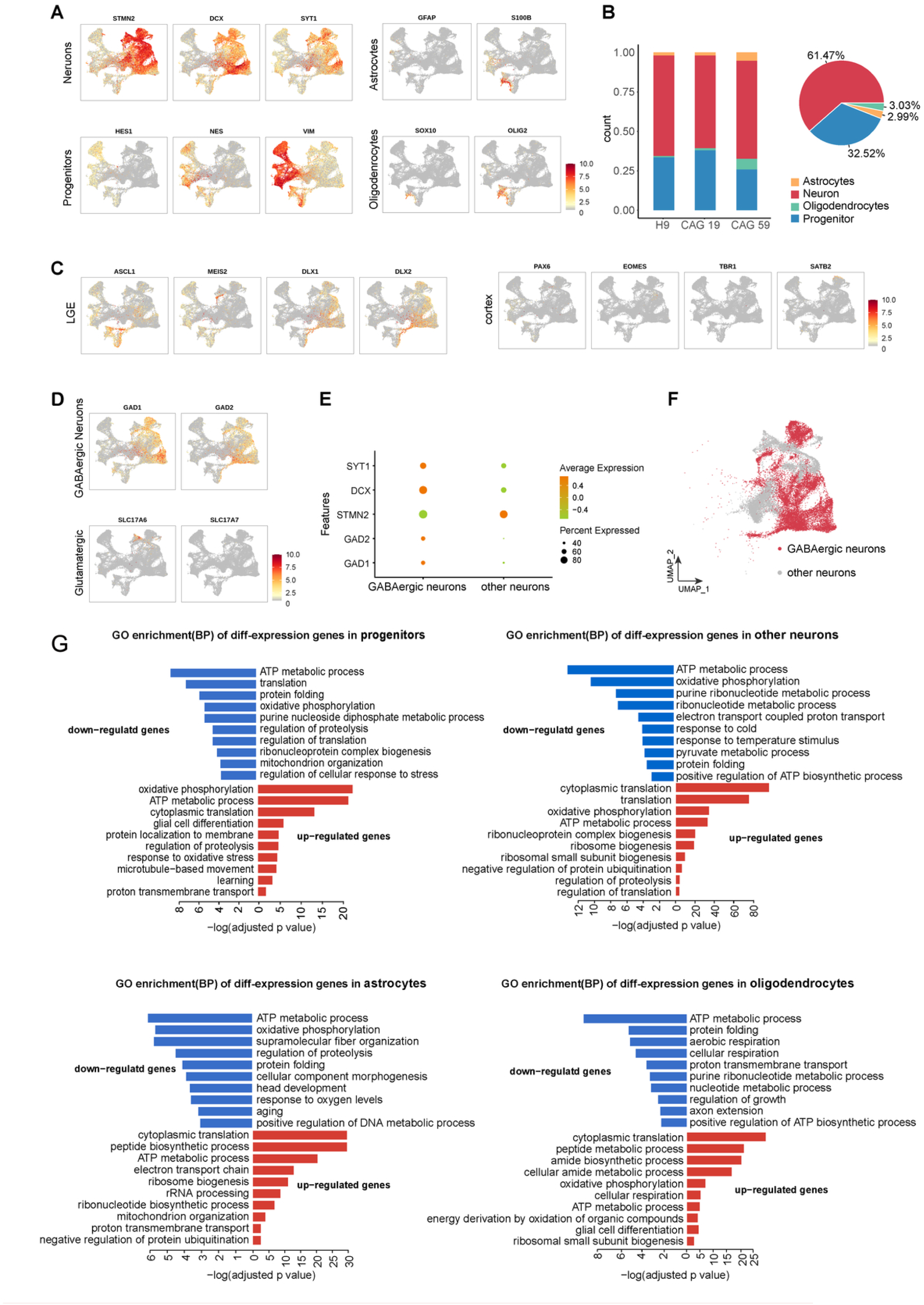
Characterizing and sorting the cellular types from scRNA-seq data of hStrOs and GO analyses of different cellular clusters in scRNA-seq data of hStrOs. **A.** Expression of known markers of progenitor, neuron, astrocyte, and oligodendrocyte. **B.** Stacked bar plots and a pie chart representing the proportion of cell types. **C, D.** Expression of known markers of LGE progenitors and cortical progenitors(C), GABAergic neurons and glutamatergic neurons (D). **E.** Dot plot displaying Expression of GABAergic neuron marker genes. **F.** UMAP plot of neurons colored by GABAergic neurons (red) and other neurons (grey). **G.** GO enrichment analysis of down-regulated (top) and up-regulated (bottom) genes in CAG59 VS CAG19/H9 progenitor, other neurons, astrocytes, and oligodendrocytes.

**Figure S8.**
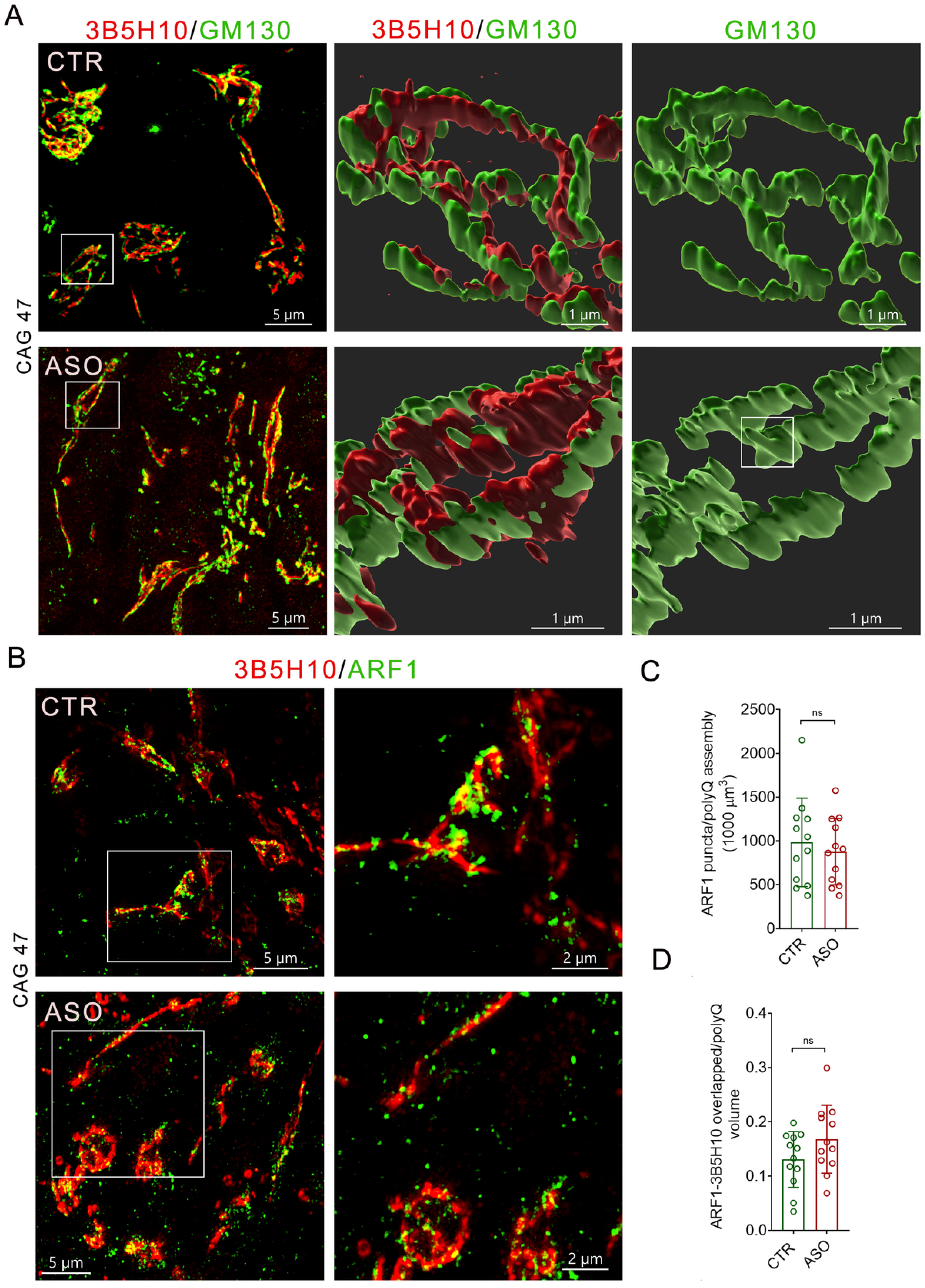
ASO treatments did not alter Golgi morphology and ARF1 scaffolding in polyQ assembly. **A.** Representative super-resolution images of polyQ assemblies stained by 3B5H10 antibody and Golgi stained by GM130 antibody in ASO-treated and control h hCOs. The middle and right panels show the spatial relationships between polyQ assemblies and the Golgi, and the tomography of the Golgi, respectively (the white rectangle, the rendering region). The renderings are done by IMARIS Surface. **B.** Representative super-resolution images of polyQ assemblies stained by 3B5H10 antibody and ARF1 puncta in ASO-treated and control (CTR) hCOs (the white rectangle, the magnified region on the left). The images are displayed as MIP. The right panels are the magnified regions in the rectangle on the left. **C.** The density of ARF1^+^ puncta in polyQ assemblies in ASO-treated and control (CTR) hCOs (HD, n=2 [CAG 47 and CAG 59]; hCOs, n=6; t-test, ns, *p*>0.05). Puncta are counted with the IMARIS surface. **D.** The overlapped volume of ARF1 puncta with polyQ assemblies in ASO-treated and control (CTR) hCOs. (HD, n=2 [CAG 47 and CAG 59]; hCOs, n=6; t-test, ns, *\*p*>0.05). PolyQ assembly and puncta volume are measured by the IMARIS surface.

## Supplementary Materials and Methods

### Primer and sequencing

A nurse collected blood from the upper arm vein, and a Blood DNA Isolation Kit extracted blood DNA. After PCR amplification, the mutant band was separated from the wild-type band on a 4% Agarose gel, and the band was then cut and extracted using a DNA Gel Extraction Kit (GenElute™, Sigma-Aldrich). The extract was sequenced by the Sanger method.

The sequencing primers:

A1: 5’-CAGAGCCCCATTCATTGCC-3’

B1: 5’-TGAGGAAGCTGAGGAGGC-3’

### Glucose starvation assays for polyQ assemblies and the Golgi

Primary fibroblasts were seeded onto 20 mm glass coverslips in 24-well plates. After 48 hours of initial culture, the standard growth medium was replaced with serum-free, low-glucose medium. To track progressive changes, cells were fixed at 24, 48, and 72 hours post-treatment and immunostained with 3B5H10 and phalloidin antibodies. For the recovery group, the serum-free, low-glucose medium was removed at the 72-hour mark and replaced with serum-free, high-glucose medium for an additional 48 hours of culture before fixation and identical immunostaining. To evaluate morphological alterations in the Golgi apparatus, a separate subset of fibroblasts was fixed 48 hours after low-glucose induction and immunostained with anti-GM130 antibodies. Fluorescent images were captured and processed using Imaris software for three-dimensional rendering. Quantification of polyQ assemblies was performed using ImageJ software.

### Onjisaponin and ASO treatments

The sequence of HuASO: 5’-CTCAGtaacattgacACCAC-3’. 5mg HuASO was synthesized by WuXi AppTec. ASOs were solubilized in 0.9% sterile saline solution or PBS. Onjisaponin F was purchased from Jindu Biotech of Shanghai and dissolved in DMSO to a concentration of 100 mM.

Brain organoids (day 100) were treated continuously for 10 days with ASOs at final concentrations of 40 µg/mL. Moreover, fibroblasts were treated continuously for 5 days with ASOs at final concentrations of 20 and 40 µg/mL. Vehicle (PBS) was used as a control. ASOs were solubilized in sterile PBS and supplemented directly into the culture medium, with media and ASO replenishment every 24 hours.

### Immunostaining and antibodies

Cells, tissues, or organoids were fixed in fresh ice-cold 4% PFA for 30 minutes and overnight, respectively. After fixation, immunostaining was performed as previously described^1^. Primary antibodies were used as follows: anti-Tuj-1 antibodies (mouse, Sigma, T8860, 1:5000; rabbit, Covance Research Products, PRB-435P, 1:5000), 3BH10 antibody (Mouse, Sigma, P1874, 1:10000), anti GABA antibodies (rabbit, Sigma, A2052, 1:5000; mouse; Sigma, A0310, 1:500), Clathrin antibody (Rabbit, Abcam, ab21679, 1:1000), EM48 (1:1000, MAB5374, Sigma-Aldrich), GM130 antibody (Rabbit, Abcam, ab52649, 1:1000), IHC-plus™ Polyclonal Rabbit anti-Human HTT / Huntingtin Antibody (C-Terminus, LS-B11011, 1:200). HTT antibody (3E10, sc-47757, 1:400), ARF1 (rabbit, 10790-1-AP, Proteintech, 1:1000), HAP40 Polyclonal Antibody (Rabbit, PA561382, Invitrogen, 1;500), CTIP2 (rat ab18465, Abcam 1:1000), PAX6 (rabbit, 901301, Biolegend, 1:1000), TBR2 (sheep, AF6166, R&D, 1:500), TBR1 (rabbit, ab31940, Abcam, 1:500), anti-MASH1 (mouse, BD, 556604,1:500), anti-DARPP32 (rabbit, Chemicon, AB1656, 1:1,000), anti-GSH2 (rabbit, Millipore, 3388451, 1:500), SATB2 (mouse, SC81376, Santa Cruz, 1:800), HOPX (rabbit, Sigma, HPA030180, 1:1000). Alexa fluor 555 labeled Wheat Germ Agglutinin (MK, MP6334).

Secondary antibodies used were as follows: Alexa Fluro 488 Donkey anti-mouse IgG (Invitrogen, Molecular Probe, A21202, 1:1000), Alexa Fluro 594 Donkey anti-mouse IgG (Invitrogen, Molecular Probe, A21203,1:1000), Alexa Fluro 594 Donkey anti-rabbit IgG(Chemicon, A21207,1:1000), Alexa Fluro 488 Donkey anti-rabbit IgG (Invitrogen, Molecular Probe, A21206,1:1000), Alexa Fluro 594 Donkey anti-goat IgG (Invitrogen, Molecular Probe, A11058,1:1000) Cy5 AffiniPure Donkey Anti-Goat IgG (H+L) (Jackson, 705-175-147,1:300), Cy5 AffiniPure Donkey Anti-Rabbit IgG (H+L) (Jackson, 711-175-152,1:300), Alexa Fluo 633 Phalloidin (Invitrogen, A22284,1:200), Mito-Tracker™ Green (Invitrogen, M7514), DAPI (Sigma, D9542), Phalloidin Alexa-633 or 488 (ThermoFisher, A12379 or A22284, 1:1000), GOLGI ID® Green assay kit (ENZO, ENZ-51028-K100).

### Golgi apparatus-selective staining and imaging in live cells

Fibroblasts were re-plated into 35 mm glass-bottomed dishes (BS-20-GJM; Biosharp). Live-cell labeling of the Golgi apparatus was performed using the GOLGI ID Green assay kit (ENZ-51028-K100; Enzo Life Sciences) following the manufacturer’s protocol. Briefly, the Assay Solution and Detection Reagent were thoroughly mixed to prepare the working staining solution. Cells were washed once with the Assay Solution, covered completely with a sufficient volume of the prepared Detection Reagent, and incubated at 37°C for 30 minutes. Following incubation, the cells were washed three times and incubated in fresh culture medium at 37°C for an additional 30 minutes prior to imaging. Time-lapse live imaging was performed by capturing sequential images from multiple regions of interest (ROIs) within the dishes at specific intervals of 30, 60, 120, and 240 minutes.

### 2D Neural Differentiation

Wild-type (WT-iPS) and Huntington’s disease (HD-iPS) induced pluripotent stem cells were differentiated into GABAergic or glutamatergic neurons using a modified protocol based on previous work^1^. Briefly, dissociated WT-iPS and HD-iPS cells were plated on Matrigel (BD) at a density of 3.5×104 cells/cm2 and maintained in E8 medium (Thermo Fisher Scientific) for 4-5 days until they reached 90% confluence. After digestion with EDTA, 9000 cells were plated in each well of U-shaped 96-well plates (SUNILON). On the first day, we maintained these cells in E8 and hNM (1:1 mixture). On the second day, the medium was replaced with hNM. From day 0 to day 6, the medium containing 2 μMol/L SB431542 (Tocris) and 0.3 μMol/L LDN193189 (BMP inhibitor, Tocris) was used for culture. These cells were plated in a well on day 7, and 200 ng/ml SHH or 0.65 μM Purmorphamine was supplied on day 10 for the differentiation of GABAergic neurons. The cells were maintained in the same medium for 2 weeks. The medium without SHH or Purmorphamine was used for differentiating glutamate neurons. At 15 days of differentiation, the rosettes were collected and cultured in a suspension condition for 25 days. On day 25, these cells were plated on a cover glass double-coated by polyonithin and laminin after digesting with accutase and then supplemented with a neural induction medium (composed of Neurobasal, N2 100×, B27 50×) (Invitrogen), supplemented with 10 ng/ml BDNF (PeproTech), 10 ng/ml GDNF (PeproTech), and 10 ng/ml IGF (PeproTech).

### Human brain organoid construction

Brain organoids were constructed according to our protocol 2. Briefly, ES/iPS colonies were dissociated into single cells using Accutase (Gibco, A1110501). 9,000 dissociated cells were plated into each well of a V-bottom ultra-low-attachment 96-well plate (Sumitomo Bakelite, MS-9096VZ). Induction media contained Human Neural Differentiation Medium and E8 media (1:1), supplemented with the two SMAD inhibitors 0.3μM LDN-193189 (STEMGENT, 040074), 2μM SB431542 (Ametek Scientific, DM-0970), and 10 μm Y-27632 (APE.BIO, A3008). On the sixth day in suspension, hCOs were transferred to neural differentiation media containing a 1:1 mixture of DMEM-F12 media and Neurobasal media supplemented with 1% (v/v) MEM-NEAA (Gibco, 11140-050), 1% (v/v) GlutaMAX(Gibco, 35050-061), 1% (v/v) N2 supplement (Gibco, 17502-048), 2% (v/v) B27 supplement (Gibco), 1% (v/v) Penicillin/Streptomycin (Gibco, 2051357). Growth factors 10ng/ml EGF (R&D Systems, 236-EG) and 20 ng/ml bFGF (R&D Systems, 233FB/CF) were used from day 6 to day 23. Growth factors, 20 ng/ml BDNF (Peprotech, AF450002) and 10 ng/ml GDNF (Peprotech, AF450010), were used from day 24 to day 45.

### Interfacing the brain organoid with HD-MEA

To facilitate robust hCO-electrode coupling, high-density microelectrode array (HD-MEA) chips (Maxlab Biosystems, Switzerland) were first functionalized. The sensing area was treated with a 0.07% (v/v) poly(ethylenimine) (PEI; Sigma-Aldrich) solution followed by a coating of laminin. For the interfacing of hCOs, we adapted the manufacturer’s protocols for brain slices. Individual hCOs were meticulously transferred to the center of the coated electrode arrays. To ensure mechanical stability and intimate contact between the basal surface of the organoid and the electrodes, a small droplet of Matrigel (Corning) was applied to secure the tissue. The chips were then lidded and incubated at 37 °C with 5% CO_2_ for 30 minutes to allow for matrix gelation. Subsequently, the culture was replenished with BrainPhys neuronal medium (StemCell Technologies) supplemented with the following reagents: 2% (v/v) NeuroCult SM1 Neuronal Supplement (StemCell Technologies), 1% (v/v) N2 Supplement-A (Gibco), 1x GlutaMAX (Gibco), 20 ng/ml recombinant human BDNF, 20 ng/ml recombinant human GDNF (PeproTech), 20 ng/ml NT-3 (PeproTech), 1 mM dibutyryl-cAMP (Sigma-Aldrich), and 1x Antibiotic-Antimycotic (Gibco). The recording medium was changed every 2–3 days.

### Electrophysiological recordings

To probe network activity in hCOs, longitudinal HD-MEA recordings were used. The spontaneous activity was recorded sequentially in blocks across the entire array (30 seconds per recording configuration, each comprising 1020 recording electrodes). Signals were sampled at 20 kHz for all recordings and saved in an HDF5 file format. Spike detection was performed using a threshold set to 5× the root-mean-square (RMS) of the noise in the band-pass-filtered signal. After the activity scan, the readout electrodes were selected depending on the requirements of each analysis (single-cell or network-related features). All HD-MEA recordings of hCOs started at least 7 days after plating. The reported electrophysiological results are based on data obtained from n=3 hCOs (CTR group), n=3 hCOs (HD group), and n=3 hCOs (HD group with ASO treatment).

### Preparation and data analysis of a single-cell RNA-sequencing library

110 days differentiated hStrOs from an HD patient (CAG 59), a healthy sibling, and H9 in suspension culture are prepared for analysis. A sterile dish with 10 mL 1× Dulbecco’s Phosphate-Buffered Saline (DPBS; Thermo Fisher, 14190144) on ice is used to transport samples after removing the residual solution. Neurosphere Dissociation Kit (Miltenyi, 130-095-943) was used to create single-cell suspensions. Dissociated cells were rinsed 1x DPBS containing 2% FBS. 0.4% Trypan blue staining (Thermo Fisher, 14190144) was used to assess viability on the Countess® II Automated Cell Counter (Thermo Fisher).

### 10x library preparation and sequencing

Beads with unique molecular identifiers (UMI) and cell barcodes were loaded close to saturation, so that each cell was paired with a bead in a Gel Beads-in-emulsion (GEM). After exposure to the cell lysis buffer, polyadenylated RNA molecules hybridized to the beads. Beads were retrieved into a single tube for reverse transcription. For cDNA synthesis, each cDNA molecule was tagged at the 5’ end (i.e., the 3’ end of a messenger RNA transcript) with a UMI and a cell label indicating its cell of origin. Briefly, 10× beads were then subjected to second-strand cDNA synthesis, adaptor ligation, and universal amplification. Sequencing libraries were prepared from randomly interrupted whole-transcriptome amplification products to enrich for the 3’ ends of transcripts linked to the cell barcode and UMI. All remaining procedures, including library construction, were performed according to the manufacturer’s standard protocol (Chromium Single Cell 3ʹ v3). A High Sensitivity DNA Chip (Agilent) on a Bioanalyzer 2100 and the Qubit High Sensitivity DNA Assay (Thermo Fisher Scientific) quantified NovaSeq6000 (Illumina) sequencing libraries (2x150 chemistry).

### Single-cell RNA-seq data processing and analysis

Cell Ranger 4.0 pipeline, with default and recommended parameters, processed the reads. Gene-Barcode matrices were generated for each sample by counting UMIs and filtering out non-cell-associated barcodes, and a gene-barcode matrix containing the barcoded cells and gene expression counts was generated. After obtaining the single-cell expression matrix, we used the Seurat (v4.0.5)^2^ R toolkit to perform normalization, clustering, dimensionality reduction, and visualization. Cells that had fewer than 200 detected genes and a percentage of mitochondrial genes lower than 10% were removed. Gene counts were then log-transformed and scaled, and 3000 shared highly variable genes were identified using the FindVariableFeatures function. We applied the RunHarmony function with the Seurat-based principal component analysis (PCA) reduction method in the Harmony (v0.1.0)^3^ package to correct batch effects and generate low-dimensional matrices. A nearest-neighbor graph was calculated using FindNeighbors and clustered using FindClusters, followed by visualization of the cell clusters using uniform manifold approximation and projection (UMAP). We used the well-known marker genes to assign these clusters to corresponding cell types. Differentially expressed genes | log2FC |> 0.25 and adjusted p-value < 0.01) among GABAergic neurons (CAG 59 vs CAG 19, CAG 59 vs H9) were identified by the Seurat function FindMarkers.

### Functional enrichment

Enrichment GO categories for significantly differentially expressed genes were identified by Metascape (https://metascape.org/gp/)^4^, which utilizes a hypergeometric test and the Benjamini-Hochberg p-value correction algorithm to identify acceptable ontology terms. Adjusted p-values of GO biological processes <0.01 were considered to be significant enrichment terms. The gene set enrichment score of Golgi vesicle transport (GO:0048193) was calculated by AUCell(v.1.14.0)^5^. The AUCell_buildRankings and AUCell_calcAUC algorithms were used for ranking model building and calculating “Area Under the Curve” (AUC) scores.

### LC-MS analysis

The patients’ iPSCs were differentiated for 50 or 100 days, and the samples were then sent to the core facility of the State Key Laboratory of Genetic Engineering for analysis. The FASP digestion was adapted for the following procedures in Microcon PL-10 filters. After a three-time buffer displacement with 8 M Urea and 50 mM ABC, pH 8.5, proteins were reduced by 10 mM DTT at 37 °C for 30 min, followed by alkylation with 30 mM iodoacetamide at 25 °C for 45 min in a light-proof condition. Digestion was performed with trypsin (enzyme/protein ratio 1:50) at 37 °C for 12 h, after which the sample was rinsed with 20% ACN and the buffer was replaced three times with digestion buffer (10 mM ABC). After digestion, the solution was filtered, the filter was rinsed twice with 15% ACN, and the filtrates were pooled and vacuum-dried. LC-MS analysis was performed using a nanoflow EASYnLC 1200 system (Thermo Fisher Scientific, Odense, Denmark) coupled to an Orbitrap Exploris 480 mass spectrometer (Thermo Fisher Scientific, Bremen, Germany). A one-column system was adopted for all analyses. A homemade C18 analytical column (75 µm i.d. × 25 cm, ReproSil-Pur 120 C18-AQ, 1.9 µm (Dr. Maisch GmbH, Germany) was used to analyze samples. The mobile phases are composed of Solution A (0.1% formic acid) and Solution B (0.1% formic acid in 80%ACN). The derivatized peptides were eluted using the following gradients: 5–8% B in 2 min, 8–44% B in 98 min, 44–60% B in 3 min, 60–100% B in 2 min, 100% B for 10 min, at a flow rate of 200 nL min. High-field asymmetric-waveform ion mobility spectrometry (FAIMS) was enabled during data acquisition, with compensation voltages set to −45 and −65 V. MS1 data were collected in the Orbitrap (60,000 resolution). Charge states 2-7 were required for MS2 analysis, and a 45 s dynamic exclusion window was used. Cycle time was set at 1.5 s. MS2 scans were performed in the Orbitrap with HCD fragmentation (isolation window 1.6; 15,000 resolution; NCE 30%).

---

**S-table-1.** The baseline characteritics of HD family members.

| No | Gender | Age | F | Onset age | CAG (mutant) | CAG (wild) | Syndrome (on sampling age) | Death | Sampling year |
| --- | --- | --- | --- | --- | --- | --- | --- | --- | --- |
| 1 | F | 46 | F3 | 42 | 47 | 19 | moderate |  | 2010 |
| 2 | M | 44 | F3 | 37 | 49 | 19 | moderate | 54 | 2010 |
| 3 | M | 21 | F4 |  |  | 19 |  |  | 2010 |
| 4 | F | 28 | F4 |  |  | 19 |  |  | 2010 |
| 5 | F | 26 | F4 |  |  | 19 |  |  | 2010 |
| 6 | M | 23 | F4 | 26 | 57 | 19 | mild |  | 2010 |
| 7 | M | 27 | F4 | 23 | 59 | 19 | mild | 36 | 2010 |
| 8 | F | 19 | F4 |  |  | 19 |  |  | 2010 |
| 9 | M | 28 | F4 | 30 | 48 | 19 | mild |  | 2010 |
| 10 | F | 26 | F4 | 32 | 47 | 19 | no syndrome |  | 2010 |
| 11 | M | 5 | F5 |  | 45 | 18 | no syndrome |  | 2010 |
| 12 | F | 39 | F4 | 30 | 55 | 19 | critical | 42 | 2010 |
| 13 | F | 42 | F4 |  |  | 19 |  |  | 2010 |
| 14 | F | 23 | F4 |  |  | 19 |  |  | 2010 |
| 15 | M | 44 | F4 | 41 | NA |  | critical |  | 2010 |
| 16 | M | 17 | F5 |  | NA |  |  |  | 2010 |
| 17 | M | 33 | F4 |  |  | 19 |  |  | 2017 |
| 18 | F | 30 | F3 |  | 45 | 19 | no syndrome |  | 2017 |
| 19 | F | 52 | F4 |  | 45 | 19 | moderate |  | 2017 |
| 20 | F | 25 | F3 |  |  | 19 |  |  | 2017 |
| 21 | M | 21 | F3 |  |  | 19 |  |  | 2017 |
| 22 | M | 52 | F2 |  | NA |  | died without syndrome |  | 2019 |
| 23 | M | 30 | F4 |  | 44 | 19 | no syndrome |  | 2019 |
| 24 | F | 54 | F3 |  | 42 | 19 | moderate |  | 2019 |
| 25 | F | 56 | F3 |  | NA |  | critical |  | 2020 |
| 26 | M | 52 | F3 |  | NA |  | healthy |  | 2020 |
| 27 | F | 28 | F4 |  | NA |  | healthy |  | 2020 |
| 28 | M | 32 | F4 |  | NA |  | healthy |  | 2020 |

F, female; M, male; NA, non-assessed; F1-4 generation

## Reference

Arrasate, M., Mitra, S., Schweitzer, E.S., Segal, M.R., and Finkbeiner, S. (2004). Inclusion body formation reduces levels of mutant huntingtin and the risk of neuronal death. Nature 431, 805–810.

Barnat, M., Capizzi, M., Aparicio, E., Boluda, S., Wennagel, D., Kacher, R., Kassem, R., Lenoir, S., Agasse, F., Braz, B.Y., et al. (2020). Huntington’s disease alters human neurodevelopment. Science 369, 787–793.

Bauerlein, F.J.B., Saha, I., Mishra, A., Kalemanov, M., Martinez-Sanchez, A., Klein, R., Dudanova, I., Hipp, M.S., Hartl, F.U., Baumeister, W., et al. (2017). In Situ Architecture and Cellular Interactions of PolyQ Inclusions. Cell 171, 179–187 e110.

Brooks, S.P., and Dunnett, S.B. (2015). Mouse Models of Huntington’s Disease. Curr Top Behav Neurosci 22, 101–133.

Caviston, J.P., Ross, J.L., Antony, S.M., Tokito, M., and Holzbaur, E.L. (2007). Huntingtin facilitates dynein/dynactin-mediated vesicle transport. Proc Natl Acad Sci U S A 104, 10045–10050.

Caviston, J.P., Zajac, A.L., Tokito, M., and Holzbaur, E.L. (2011). Huntingtin coordinates the dynein-mediated dynamic positioning of endosomes and lysosomes. Mol Biol Cell 22, 478–492.

Chen, X., Saiyin, H., Liu, Y., Wang, Y., Li, X., Ji, R., and Ma, L. (2022). Human striatal organoids derived from pluripotent stem cells recapitulate striatal development and compartments. Plos Biol 20, e3001868.

Costa, V., Giacomello, M., Hudec, R., Lopreiato, R., Ermak, G., Lim, D., Malorni, W., Davies, K.J., Carafoli, E., and Scorrano, L. (2010). Mitochondrial fission and cristae disruption increase the response of cell models of Huntington’s disease to apoptotic stimuli. EMBO Mol Med 2, 490–503.

D’Souza-Schorey, C., and Chavrier, P. (2006). ARF proteins: roles in membrane traffic and beyond. Nat Rev Mol Cell Biol 7, 347–358.

del Toro, D., Alberch, J., Lazaro-Dieguez, F., Martin-Ibanez, R., Xifro, X., Egea, G., and Canals, J.M. (2009). Mutant huntingtin impairs post-Golgi trafficking to lysosomes by delocalizing optineurin/Rab8 complex from the Golgi apparatus. Mol Biol Cell 20, 1478–1492.

del Toro, D., Canals, J.M., Gines, S., Kojima, M., Egea, G., and Alberch, J. (2006). Mutant huntingtin impairs the post-Golgi trafficking of brain-derived neurotrophic factor but not its Val66Met polymorphism. J Neurosci 26, 12748–12757.

DiFiglia, M., Sapp, E., Chase, K., Schwarz, C., Meloni, A., Young, C., Martin, E., Vonsattel, J.P., Carraway, R., Reeves, S.A., et al. (1995). Huntingtin is a cytoplasmic protein associated with vesicles in human and rat brain neurons. Neuron 14, 1075–1081.

DiFiglia, M., Sapp, E., Chase, K.O., Davies, S.W., Bates, G.P., Vonsattel, J.P., and Aronin, N. (1997). Aggregation of huntingtin in neuronal intranuclear inclusions and dystrophic neurites in brain. Science 277, 1990–1993.

Dougan, L., Li, J., Badilla, C.L., Berne, B.J., and Fernandez, J.M. (2009). Single homopolypeptide chains collapse into mechanically rigid conformations. Proc Natl Acad Sci U S A 106, 12605–12610.

Dragatsis, I., Efstratiadis, A., and Zeitlin, S. (1998). Mouse mutant embryos lacking huntingtin are rescued from lethality by wild-type extraembryonic tissues. Development 125, 1529–1539.

Duyao, M.P., Auerbach, A.B., Ryan, A., Persichetti, F., Barnes, G.T., McNeil, S.M., Ge, P., Vonsattel, J.P., Gusella, J.F., Joyner, A.L., et al. (1995). Inactivation of the mouse Huntington’s disease gene homolog Hdh. Science 269, 407–410.

El Ghouzzi, V., and Boncompain, G. (2022). Golgipathies reveal the critical role of the sorting machinery in brain and skeletal development. Nat Commun 13, 7397.

Fasano, G., Muto, V., Radio, F.C., Venditti, M., Mosaddeghzadeh, N., Coppola, S., Paradisi, G., Zara, E., Bazgir, F., Ziegler, A., et al. (2022). Dominant ARF3 variants disrupt Golgi integrity and cause a neurodevelopmental disorder recapitulated in zebrafish. Nat Commun 13, 6841.

Ferrante, R.J., Gutekunst, C.A., Persichetti, F., McNeil, S.M., Kowall, N.W., Gusella, J.F., MacDonald, M.E., Beal, M.F., and Hersch, S.M. (1997). Heterogeneous topographic and cellular distribution of huntingtin expression in the normal human neostriatum. J Neurosci 17, 3052–3063.

Grima, J.C., Daigle, J.G., Arbez, N., Cunningham, K.C., Zhang, K., Ochaba, J., Geater, C., Morozko, E., Stocksdale, J., Glatzer, J.C., et al. (2017). Mutant Huntingtin Disrupts the Nuclear Pore Complex. Neuron 94, 93–107 e106.

Guo, Q., Bin, H., Cheng, J., Seefelder, M., Engler, T., Pfeifer, G., Oeckl, P., Otto, M., Moser, F., Maurer, M., et al. (2018). The cryo-electron microscopy structure of huntingtin. Nature 555, 117–120.

Gutekunst, C.A., Li, S.H., Yi, H., Mulroy, J.S., Kuemmerle, S., Jones, R., Rye, D., Ferrante, R.J., Hersch, S.M., and Li, X.J. (1999). Nuclear and neuropil aggregates in Huntington’s disease: relationship to neuropathology. J Neurosci 19, 2522–2534.

Harding, R.J., Deme, J.C., Hevler, J.F., Tamara, S., Lemak, A., Cantle, J.P., Szewczyk, M.M., Begeja, N., Goss, S., Zuo, X., et al. (2021). Huntingtin structure is orchestrated by HAP40 and shows a polyglutamine expansion-specific interaction with exon 1. Commun Biol 4, 1374.

Hatters, D.M., Hannan, A.J., and Springer Science+Business Media (2013). Tandem repeats in genes, proteins, and disease : methods and protocols (New York: Humana Press).

Isas, J.M., Langen, A., Isas, M.C., Pandey, N.K., and Siemer, A.B. (2017). Formation and Structure of Wild Type Huntingtin Exon-1 Fibrils. Biochemistry-Us 56, 3579–3586.

Jimenez-Sanchez, M., Licitra, F., Underwood, B.R., and Rubinsztein, D.C. (2017). Huntington’s Disease: Mechanisms of Pathogenesis and Therapeutic Strategies. Cold Spring Harb Perspect Med 7.

Kernich, C.A. (2002). Huntington’s disease. Neurologist 8, 207–208.

Kim, J., Moody, J.P., Edgerly, C.K., Bordiuk, O.L., Cormier, K., Smith, K., Beal, M.F., and Ferrante, R.J. (2010). Mitochondrial loss, dysfunction and altered dynamics in Huntington’s disease. Hum Mol Genet 19, 3919–3935.

Kim, M., Lee, H.S., LaForet, G., McIntyre, C., Martin, E.J., Chang, P., Kim, T.W., Williams, M., Reddy, P.H., Tagle, D., et al. (1999). Mutant huntingtin expression in clonal striatal cells: dissociation of inclusion formation and neuronal survival by caspase inhibition. J Neurosci 19, 964–973.

Klumperman, J. (2011). Architecture of the mammalian Golgi. Cold Spring Harb Perspect Biol 3.

Kordasiewicz, H.B., Stanek, L.M., Wancewicz, E.V., Mazur, C., McAlonis, M.M., Pytel, K.A., Artates, J.W., Weiss, A., Cheng, S.H., Shihabuddin, L.S., et al. (2012). Sustained therapeutic reversal of Huntington’s disease by transient repression of huntingtin synthesis. Neuron 74, 1031–1044.

Lajoie, P., and Snapp, E.L. (2010). Formation and toxicity of soluble polyglutamine oligomers in living cells. PLoS One 5, e15245.

Lebon, S., Bruneel, A., Drunat, S., Albert, A., Csaba, Z., Elmaleh, M., Ntorkou, A., Tenier, Y., Fenaille, F., Gressens, P., et al. (2025). A biallelic variant in GORASP1 causes a novel Golgipathy with glycosylation and mitotic defects. Life Sci Alliance 8.

Lee, H., Fenster, R.J., Pineda, S.S., Gibbs, W.S., Mohammadi, S., Davila-Velderrain, J., Garcia, F.J., Therrien, M., Novis, H.S., Gao, F., et al. (2020). Cell Type-Specific Transcriptomics Reveals that Mutant Huntingtin Leads to Mitochondrial RNA Release and Neuronal Innate Immune Activation. Neuron 107, 891-+.

Legleiter, J., Mitchell, E., Lotz, G.P., Sapp, E., Ng, C., DiFiglia, M., Thompson, L.M., and Muchowski, P.J. (2010). Mutant huntingtin fragments form oligomers in a polyglutamine length-dependent manner in vitro and in vivo. J Biol Chem 285, 14777–14790.

Liu, C., Mei, M., Li, Q., Roboti, P., Pang, Q., Ying, Z., Gao, F., Lowe, M., and Bao, S. (2017). Loss of the golgin GM130 causes Golgi disruption, Purkinje neuron loss, and ataxia in mice. Proc Natl Acad Sci U S A 114, 346–351.

Liu, J.Y., Huang, Y., Li, T., Jiang, Z., Zeng, L.W., and Hu, Z.P. (2021). The role of the Golgi apparatus in disease (Review). Int J Mol Med 47.

Liu, Y., Chen, X., Ma, Y., Song, C., Ma, J., Chen, C., Su, J., Ma, L., and Saiyin, H. (2024). Endogenous mutant Huntingtin alters the corticogenesis via lowering Golgi recruiting ARF1 in cortical organoid. Mol Psychiatry.

Machiela, E., and Southwell, A.L. (2020). Biological Aging and the Cellular Pathogenesis of Huntington’s Disease. J Huntingtons Dis 9, 115–128.

Martin, D.D., Ladha, S., Ehrnhoefer, D.E., and Hayden, M.R. (2015). Autophagy in Huntington disease and huntingtin in autophagy. Trends Neurosci 38, 26–35.

Mehta, S.R., Tom, C.M., Wang, Y., Bresee, C., Rushton, D., Mathkar, P.P., Tang, J., and Mattis, V.B. (2018). Human Huntington’s Disease iPSC-Derived Cortical Neurons Display Altered Transcriptomics, Morphology, and Maturation. Cell Rep 25, 1081–1096 e1086.

Mier, P., and Andrade-Navarro, M.A. (2021). Between Interactions and Aggregates: The PolyQ Balance. Genome Biol Evol 13.

Miller, J., Arrasate, M., Brooks, E., Libeu, C.P., Legleiter, J., Hatters, D., Curtis, J., Cheung, K., Krishnan, P., Mitra, S., et al. (2011). Identifying polyglutamine protein species in situ that best predict neurodegeneration. Nat Chem Biol 7, 925–934.

Milnerwood, A.J., Cummings, D.M., Dallerac, G.M., Brown, J.Y., Vatsavayai, S.C., Hirst, M.C., Rezaie, P., and Murphy, K.P. (2006). Early development of aberrant synaptic plasticity in a mouse model of Huntington’s disease. Hum Mol Genet 15, 1690–1703.

Milnerwood, A.J., Gladding, C.M., Pouladi, M.A., Kaufman, A.M., Hines, R.M., Boyd, J.D., Ko, R.W., Vasuta, O.C., Graham, R.K., Hayden, M.R., et al. (2010). Early increase in extrasynaptic NMDA receptor signaling and expression contributes to phenotype onset in Huntington’s disease mice. Neuron 65, 178–190.

Morlot, S., and Roux, A. (2013). Mechanics of dynamin-mediated membrane fission. Annu Rev Biophys 42, 629–649.

Mukai, H., Isagawa, T., Goyama, E., Tanaka, S., Bence, N.F., Tamura, A., Ono, Y., and Kopito, R.R. (2005). Formation of morphologically similar globular aggregates from diverse aggregation-prone proteins in mammalian cells. Proc Natl Acad Sci U S A 102, 10887–10892.

Nagai, T., Suzuki, Y., Kiyohara, H., Susa, E., Kato, T., Nagamine, T., Hagiwara, Y., Tamura, S., Yabe, T., Aizawa, C., et al. (2001). Onjisaponins, from the root of Polygala tenuifolia Willdenow, as effective adjuvants for nasal influenza and diphtheria-pertussis-tetanus vaccines. Vaccine 19, 4824–4834.

Olenick, M.A., and Holzbaur, E.L.F. (2019). Dynein activators and adaptors at a glance. J Cell Sci 132.

Owens, G.E., New, D.M., West, A.P., and Bjorkman, P.J. (2015). Anti-PolyQ Antibodies Recognize a Short PolyQ Stretch in Both Normal and Mutant Huntingtin Exon 1. Journal of Molecular Biology 427, 2507–2519.

Passemard, S., Perez, F., Colin-Lemesre, E., Rasika, S., Gressens, P., and El Ghouzzi, V. (2017). Golgi trafficking defects in postnatal microcephaly: The evidence for “Golgipathies”. Prog Neurobiol 153, 46–63.

Peters, P.J., Ning, K., Palacios, F., Boshans, R.L., Kazantsev, A., Thompson, L.M., Woodman, B., Bates, G.P., and D’Souza-Schorey, C. (2002). Arfaptin 2 regulates the aggregation of mutant huntingtin protein. Nat Cell Biol 4, 240–245.

Poirier, M.A., Li, H.L., Macosko, J., Cai, S.W., Amzel, M., and Ross, C.A. (2002). Huntingtin spheroids and protofibrils as precursors in polyglutamine fibrilization. Journal of Biological Chemistry 277, 41032–41037.

Rao, S., Kirschen, G.W., Szczurkowska, J., Di Antonio, A., Wang, J., Ge, S., and Shelly, M. (2018). Repositioning of Somatic Golgi Apparatus Is Essential for the Dendritic Establishment of Adult-Born Hippocampal Neurons. J Neurosci 38, 631–647.

Ravichandran, Y., Goud, B., and Manneville, J.B. (2020). The Golgi apparatus and cell polarity: Roles of the cytoskeleton, the Golgi matrix, and Golgi membranes. Curr Opin Cell Biol 62, 104–113.

Sakamoto, M., Sasaki, K., Sugie, A., Nitta, Y., Kimura, T., Gursoy, S., Cinleti, T., Iai, M., Sengoku, T., Ogata, K., et al. (2021). De novo ARF3 variants cause neurodevelopmental disorder with brain abnormality. Hum Mol Genet 31, 69–81.

Saudou, F., Finkbeiner, S., Devys, D., and Greenberg, M.E. (1998). Huntingtin acts in the nucleus to induce apoptosis but death does not correlate with the formation of intranuclear inclusions. Cell 95, 55–66.

Saudou, F., and Humbert, S. (2016). The Biology of Huntingtin. Neuron 89, 910–926.

Scherzinger, E., Sittler, A., Schweiger, K., Heiser, V., Lurz, R., Hasenbank, R., Bates, G.P., Lehrach, H., and Wanker, E.E. (1999). Self-assembly of polyglutamine-containing huntingtin fragments into amyloid-like fibrils: implications for Huntington’s disease pathology. Proc Natl Acad Sci U S A 96, 4604–4609.

Shirasaki, D.I., Greiner, E.R., Al-Ramahi, I., Gray, M., Boontheung, P., Geschwind, D.H., Botas, J., Coppola, G., Horvath, S., Loo, J.A., et al. (2012). Network organization of the huntingtin proteomic interactome in mammalian brain. Neuron 75, 41–57.

Silvestroni, A., Faull, R.L., Strand, A.D., and Moller, T. (2009). Distinct neuroinflammatory profile in post-mortem human Huntington’s disease. Neuroreport 20, 1098–1103.

Sivaramakrishnan, S., Spink, B.J., Sim, A.Y., Doniach, S., and Spudich, J.A. (2008). Dynamic charge interactions create surprising rigidity in the ER/K alpha-helical protein motif. Proc Natl Acad Sci U S A 105, 13356–13361.

Stanley, P. (2011). Golgi glycosylation. Cold Spring Harb Perspect Biol 3.

Tabrizi, S.J., Flower, M.D., Ross, C.A., and Wild, E.J. (2020). Huntington disease: new insights into molecular pathogenesis and therapeutic opportunities. Nat Rev Neurol 16, 529–546.

Takahashi, T., Kikuchi, S., Katada, S., Nagai, Y., Nishizawa, M., and Onodera, O. (2008). Soluble polyglutamine oligomers formed prior to inclusion body formation are cytotoxic. Hum Mol Genet 17, 345–356.

Tang, C.C., Feigin, A., Ma, Y., Habeck, C., Paulsen, J.S., Leenders, K.L., Teune, L.K., van Oostrom, J.C., Guttman, M., Dhawan, V., et al. (2013). Metabolic network as a progression biomarker of premanifest Huntington’s disease. J Clin Invest 123, 4076–4088.

Tousley, A., Iuliano, M., Weisman, E., Sapp, E., Richardson, H., Vodicka, P., Alexander, J., Aronin, N., DiFiglia, M., and Kegel-Gleason, K.B. (2019). Huntingtin associates with the actin cytoskeleton and alpha-actinin isoforms to influence stimulus dependent morphology changes. Plos One 14.

Truant, R., Atwal, R.S., Desmond, C., Munsie, L., and Tran, T. (2008). Huntington’s disease: revisiting the aggregation hypothesis in polyglutamine neurodegenerative diseases. FEBS J 275, 4252–4262.

Villavicencio Gonzalez, E., and Zoghbi, H.Y. (2026). Pathogenesis of polyglutamine diseases: Piecing together a complex molecular puzzle. J Exp Med 223.

Walker, F.O. (2007a). Huntington’s disease. Lancet 369, 218–228.

Walker, F.O. (2007b). Huntington’s disease. Semin Neurol 27, 143–150.

Wei, J.H., and Seemann, J. (2009). Mitotic division of the mammalian Golgi apparatus. Semin Cell Dev Biol 20, 810–816.

Wu, L.L., and Zhou, X.F. (2009). Huntingtin associated protein 1 and its functions. Cell Adh Migr 3, 71–76.

Yadav, S., and Linstedt, A.D. (2011). Golgi positioning. Cold Spring Harb Perspect Biol 3.

Yang, H., Yang, S., Jing, L., Huang, L., Chen, L., Zhao, X., Yang, W., Pan, Y., Yin, P., Qin, Z.S., et al. (2020). Truncation of mutant huntingtin in knock-in mice demonstrates exon1 huntingtin is a key pathogenic form. Nat Commun 11, 2582.

Zeitlin, S., Liu, J.P., Chapman, D.L., Papaioannou, V.E., and Efstratiadis, A. (1995). Increased apoptosis and early embryonic lethality in mice nullizygous for the Huntington’s disease gene homologue. Nat Genet 11, 155–163.

## Reference

1 Ma, L. et al. Human embryonic stem cell-derived GABA neurons correct locomotion deficits in quinolinic acid-lesioned mice. Cell Stem Cell 10, 455–464, doi:10.1016/j.stem.2012.01.021 (2012).

2 Butler, A., Hoffman, P., Smibert, P., Papalexi, E. & Satija, R. Integrating single-cell transcriptomic data across different conditions, technologies, and species. Nature Biotechnology 36, 411–+, doi:10.1038/nbt.4096 (2018).

3 Korsunsky, I. et al. Fast, sensitive and accurate integration of single-cell data with Harmony. Nature Methods 16, 1289–+, doi:10.1038/s41592-019-0619-0 (2019).

4 Zhou, Y. Y., et al. Metascape provides a biologist-oriented resource for the analysis of systems-level datasets. Nat Commun 10, doi:ARTN 1523 10.1038/s41467-019-09234-6 (2019).

5 van de Sande, B. et al. A scalable SCENIC workflow for single-cell gene regulatory network analysis. Nature Protocols 15, 2247–2276, doi:10.1038/s41596-020-0336-2 (2020).

